# Reciprocal stabilisation of glycine receptors and gephyrin scaffold proteins at inhibitory synapses

**DOI:** 10.1101/2020.12.14.422654

**Authors:** Thomas Chapdelaine, Vincent Hakim, Antoine Triller, Jonas Ranft, Christian G Specht

## Abstract

Postsynaptic scaffold proteins immobilise neurotransmitter receptors in the synaptic membrane opposite to presynaptic vesicle release sites, thus ensuring efficient synaptic transmission. At inhibitory synapses in the spinal cord, the main scaffold protein gephyrin assembles in dense molecule clusters that provide binding sites for glycine receptors (GlyRs). Gephyrin and GlyRs can also interact outside of synapses where they form receptor-scaffold complexes. While several models for the formation of postsynaptic scaffold domains in the presence of receptor-scaffold interactions have been advanced, a clear picture of the coupled dynamics of receptors and scaffold proteins at synapses is lacking.

To characterise the GlyR and gephyrin dynamics at inhibitory synapses we performed fluorescence time-lapse imaging after photoconversion in order to directly visualise the exchange kinetics of recombinant Dendra2-gephyrin in cultured spinal cord neurons. Immuno-immobilisation of endogenous GlyRs with specific antibodies abolished their lateral diffusion in the plasma membrane, as judged by the lack of fluorescence recovery after photobleaching. Moreover, the crosslinking of GlyRs significantly reduced the exchange of Dendra2-gephyrin compared to control conditions, suggesting that the kinetics of the synaptic gephyrin pool is strongly dependent on GlyR-gephyrin interactions. We did not observe any change in the total synaptic gephyrin levels after GlyR crosslinking, however, indicating that the number of gephyrin molecules at synapses is not primarily dependent on the exchange of GlyR-gephyrin complexes.

We further show that our experimental data can be quantitatively accounted for by a model of receptor-scaffold dynamics that includes a tightly interacting receptor-scaffold domain, as well as more loosely bound receptor and scaffold populations that exchange with extrasynaptic pools. The model can make predictions for single molecule data such as typical dwell times of synaptic proteins. Taken together, our data demonstrate the reciprocal stabilisation of GlyRs and gephyrin at inhibitory synapses and provide a quantitative understanding of their dynamic organisation.

**Statement of significance:** The efficiency of signal transmission between neurons depends strongly on the number of available neurotransmitter receptors in the postsynaptic membrane. Postsynaptic scaffold proteins provide binding sites for receptors, thus setting the gain of synaptic transmission. However, the importance of receptor-scaffold interactions for the stability of the postsynaptic scaffold itself has received relatively little attention. Using time-lapse imaging of glycine receptors and gephyrin scaffolds at inhibitory synapses in spinal cord neurons together with biophysical modelling, we show that receptor mobility controls the exchange but not the total number of gephyrin molecules at the synapse, and predict that glycine receptors and gephyrin scaffolds dynamically organise into different subpopulations with varying degrees of reciprocal stabilisation.

## Introduction

The postsynaptic scaffold at inhibitory synapses is characterised by the presence of dense clusters of gephyrin molecules that provide binding sites for inhibitory glycine receptors (GlyR) and GABA type A receptors (GABA_A_Rs), as well as other synaptic components such as collybistin and neuroligin-2 (reviewed in (1)). Gephyrin has a particularly strong interaction with the intracellular domain of the β-subunit of the GlyR with a K_D_ in the nanomolar range (2-5)). The presence of the β-subunit is therefore essential to anchor the pentameric GlyR complex in the postsynaptic membrane (6, 7). Gephyrin is also involved in the forward trafficking of GlyRs towards the plasma membrane (8, 9), where it remains associated with the receptor due to the high affinity of the GlyRβ-gephyrin interaction (7, 10).

In addition to receptor-scaffold interactions, the stability of inhibitory synapses is also dependent on scaffold-scaffold interactions. The basic unit of soluble gephyrin is a trimer, formed by homomeric interactions between the N-terminal domains (G-domains) of gephyrin (11, 12). Furthermore, the C-terminal domains of gephyrin (E-domains) can under certain conditions form dimers (2, 13, 14) that are thought to be required for synaptic clustering (15) (discussed in (1)).

The different molecular states of GlyRs and gephyrin are engaged in a dynamic equilibrium that can be largely accounted for by receptor-gephyrin and gephyrin-gephyrin interactions (10). In support of this view, expression of gephyrin and GlyRs in non-neuronal cells is sufficient to drive the spontaneous formation of membrane-associated gephyrin aggregates that resemble postsynaptic domains (e.g. (16, 17)). Several models have been put forward to explain the formation of stable gephyrin domains arising from receptor-scaffold interactions (17-20).

Based on thermodynamic considerations, Sekimoto and Triller proposed a mechanism of phase separation between a condensed domain (phase) at synapses and a delocalised phase with lower receptor and gephyrin concentrations in the extrasynaptic space (18). Haselwandter and colleagues treated the GlyR and gephyrin populations as a reaction-diffusion system and proposed that postsynaptic domains are formed by a Turing-like instability (17, 21). More recently, we hypothesised that gephyrin domains are in a nonequilibrium stationary state where the desorption of synaptic gephyrin proteins into the cytoplasm is balanced by the capture of diffusing GlyR-gephyrin complexes (19, 20). In this project, we set out to put these different ideas to the test by directly measuring the exchange dynamics of GlyRs and gephyrin at synapses using population measurements based on photoconversion and time-lapse imaging in cultured spinal cord neurons.

## Methods

### Neuron culture and lentivirus infection

Primary spinal cord neurons of Sprague-Dawley rat embryos at embryonic stage E14 were cultured as described previously (22). Neurons were plated and grown on 18 mm diameter glass coverslips in Neurobasal medium containing complement B27, 2 mM glutamine, 5 U/ml penicillin and 5 μg/ml streptomycin at 37°C and 5% CO_2_. Half of the culture medium was replaced twice a week with BrainPhys medium containing SM1 and antibiotics. Cultures were infected at day in vitro 3 or 4 (DIV3-4) with lentivirus (15 μl per coverslip) driving the expression of Dendra2-gephyrin or Dendra2-GlyRα1 (see Supplementary Methods), and used for experiments between DIV13 and DIV16.

### Antibody crosslinking and live imaging

Before each experiment, the coverslips were rinsed in warm Tyrode solution (120 mM NaCl, 2.5 mM KCl, 2 mM CaCl_2_, 2 mM MgCl_2_, 25 mM glucose, 5 mM pyruvate and 25 mM HEPES, pH 7.4) and placed on a heating plate at 37°C. GlyRs were immuno-immobilised by incubating the cultured spinal cord neurons with primary rabbit anti-GlyRα1 antibody (custom-made, 1:100 dilution in Tyrode solution) for 10 minutes, rinsed twice, and incubated for another 10 minutes with Alexa Fluor 647 or Alexa Fluor 488 conjugated secondary antibodies (A647-coupled donkey anti-rabbit, A488 goat anti-rabbit, Jackson, 1:100). Coverslips were rinsed again, mounted in an imaging chamber on the microscope stage, and imaged for up to one hour (typically 35-40 min) at 37°C in Tyrode solution. Temperature and humidity were maintained using a H301-T-UNIT-BL-PLUS temperature control unit (Okolab). Whenever direct control experiments were carried out (data in Fig. 3 and S5), all coverslips were treated in the same way, using Tyrode solution without antibodies in the control condition.

### Photoconversion and time-lapse image acquisition

To determine the most suitable fluorophore for the photoconversion experiments, we compared the behaviour of different photoconvertible fluorescent proteins (Dendra2, mEos2, mEos4b) in COS-7 cells (Fig. S1). We noticed a strong increase of the fluorescence intensity of non-converted mEos2 and mEos4b in response to low intensity UV illumination. This photochromism of the Eos fluorophores introduces a non-linearity in the intensity measurements that complicates data analysis. We therefore chose Dendra2 for our experiments, since this fluorophore was the least affected by photochromic effects.

Fluorescence recovery after photobleaching (FRAP) and fluorescence decay after photoconversion (FDAP) was carried out in cultured spinal cord neurons expressing Dendra2-tagged gephyrin or GlyRs. Images were acquired on an inverted Nikon Eclipse Ti microscope equipped with a perfect focus system (Nikon), a 100x oil-immersion objective (Nikon Apochromat, NA 1.49), a 1.5x magnifying lens, a module for focusing the laser beam (Ti-FRAP, Nikon, spot size ~1 μm^2^), and an EMCCD camera (Andor iXon Ultra, 512 x 512 pixels). For wide-field imaging, neurons were illuminated with a Solis-1C LED lamp (Thorlabs, set at 1000 mA) using specific excitation band-pass filters (485/20 nm, 560/25 nm, 650/13 nm, Semrock), a multi-band dichroic mirror (410/504/582/669 nm), and the appropriate emission filters (440/40 nm, 525/30 nm, 607/36 nm, 684/24 nm).

Photoconversion of Dendra2-gephyrin and Dendra2-GlyRα1 was done with a 405 nm laser (Obis Coherent, 120 mW). Alternatively, GlyRs that were immuno-immobilised and labelled with A488 (data in Fig. 2) were photobleached using a 488 nm laser (Obis Coherent, 150 mW). The intensity of the laser pulse was controlled by an acousto-optic tunable filter (AOTF) and introduced through an optical fibre via the upper filter turret of the microscope using a multi-band dichroic mirror (405/488/543/635 nm) that was positioned in the light path only during laser illumination.

Photoconversion and acquisition parameters for time-lapse imaging were set in fixed neurons expressing Dendra2-gephyrin (Fig. S2). Images were acquired with NIS Elements software (Nikon) according to the following sequence: a single image was taken in the far red channel, followed by three pairs of images every 10 seconds in the red and the green channels (baseline before FRAP/FDAP). Then, the 405 nm or 488 nm laser pulse was applied, after which a further sixteen images were taken at regular intervals (10 s in fixed neurons) in the red and the green channels, followed by one final image in the far red channel. In live cell experiments, time-lapse images in the green and the red channels were acquired every 2 minutes or every 15 seconds with 200 ms exposure using a neutral density filter (ND 8) to obtain the best compromise between image quality, temporal resolution and bleaching. Photobleaching in fixed samples was below 0.4% per acquired image in the green channel prior to FRAP/FDAP, and no further bleaching was detected after photoconversion in both channels throughout the recording (Fig. S2). The laser intensity was adjusted to maximise the rate of photoconversion. Application of a single 405 nm pulse (5% of the maximal laser output, 500 ms) reduced the average intensity of the green fluorescence in the targeted area (~1 μm^2^ spot) close to background levels (approx. 50% of the baseline before FRAP/FDAP), while producing large gains in red fluorescence (about 300 a.u. above background in fixed samples). In the GlyR crosslinking experiments with A488-conjugated secondary antibodies, the dyes were bleached with a single pulse of a 488 nm laser (20% intensity, 1 s).

### Image processing and data analysis

FRAP/FDAP image stacks were separated by channel. Seven areas of 9 x 9 pixels (106 nm pixel size, i.e. squares of ~1 μm^2^) were chosen in the green channel: one covering the synapse that was targeted by photoconversion (FRAP/FDAP), three synaptic puncta close to the centre of the image (near controls) and three synaptic puncta far from the centre (far controls). Two additional zones of variable dimensions were defined: one on the soma or on a segment of dendrite as a measure of the diffuse level of fluorescence of the neuron (background), and another outside the cell to determine the non-specific fluorescence (offset). The selected points were tracked automatically using Openview software (Noam Ziv, Technion, Israel Institute of Technology), applying a manual correction if the position of the spot was lost after photoconversion. The average fluorescence intensity of the tracked spots (9 x 9 pixels) was measured in each channel (Fig. S2).

#### Data curation of the live Dendra2-gephyrin experiments

In order to homogenize the distribution of the initial fluorescence of the selected FRAP/FDAP spots between control and immuno-immobilised conditions, we excluded outliers in the control condition for which the average intensity in the green channel before FRAP/FDAP exceeded 3000 a.u. (the maximum pre-FRAP/FDAP intensity observed in the immuno-immobilised condition was 2490 a.u.). This is equivalent to requiring that intensities lie within 3.5 median absolute deviations (MAD) of the median for the control condition, where 5 of 57 recordings were rejected based on this criterion. We furthermore excluded recordings in which the application of the FRAP/FDAP laser pulse did not produce a significant drop in fluorescence in the green channel. Specifically, we required that the intensity drop relative to the average pre-FRAP/FDAP intensity exceeded the baseline fluctuations of fluorescence intensity by a factor of four, where the size of the fluctuations was quantified by the standard deviation of the intensity measured in the three images taken before the pulse. In the control (immuno-immobilised) condition, this led to the exclusion of another four (two) recordings.

#### FRAP data analysis

In a pilot experiment with Dendra2-gephyrin in living neurons, we observed that low intensity 405 nm light triggered a slight increase of the green fluorescence intensity (Fig. S3), a behaviour that had not been seen in fixed samples (Fig. S1, S2). This photochromism was factored out by normalising the recovery data with the near control puncta. We applied the following normalisation procedure: First, the intensity *I*_raw_(*t*) of each FRAP spot was corrected by a multiplicative factor *n*(*t*) that accounts for the time-dependent overactivation of the near control puncta, and was determined as the average intensity of all near control puncta at time *t* divided by the average intensity of all near control puncta in the three images taken before FRAP. The corrected intensity *I_corr_*(*t*) = *I*_raw_(*t*)/*n*(*t*) was then normalised and rescaled relative to its pre-FRAP average *I*_pre_ and immediate post-FRAP value *I*_0_ according to *I*_norm_(*t*) = (*I*_corr_(*t*) = *I*_0_)/(*I*_pre_ − *I*_0_). We then used a two-parameter exponential fit to characterise the observed FRAP dynamics (see Statistics and fitting).

#### FDAP data analysis

We normalised and rescaled the fluorescence intensity *I*(*t*) in the red channel to its pre-FDAP average *I*_pre_ and immediate post-FDAP value *I*_0_ according to *I*_norm_(*t*) = (*I*(*t*) − *I*_pre_)/(*I*_0_ − *I*_pre_). The intensity of the converted spot in the first image after the 405 nm pulse was about 15% higher than in all subsequent images, both in fixed and live samples, a likely consequence of 560/25 nm excitation of newly converted Dendra2 fluorophores. We accounted for this overactivation by including an additional offset *f*_off_ in the exponential fit of the FDAP dynamics (see below).

#### Combination of FRAP and FDAP data

In the stationary state and in the absence of imaging artefacts, nonlinearities etc., the average, normalised FRAP signal should follow the same dynamics as the average, normalised FDAP signal, with FDAP_theoretical_(*t*) = 1 − FRAP_theoretical_(*t*). To make use of both FRAP and FDAP data in our theoretical model, we therefore fitted our model (Fig. 5) to averages of the FRAP and FDAP traces, where we corrected the FDAP data for the overactivation at *t* = 0 min using the offset that best fitted the experimental data (see below): FRAP_data,combined_(*t*) = [FRAP_data_(*t*) + 1 − FDAP_data,corr_(*t*)]/2, with FDAP_data,corr_(*t*) = FDAP_data_(*t*)/(1 − *f*_off_).

### Statistics and fitting

Data are given in mean ± standard deviation (SD) or standard error of the mean (SEM) as indicated. Pairwise comparison of intensity data of synaptic puncta (Fig. 4) was done using a non-parametric Mann-Whitney U-test (two-tailed).

Fits were performed in Python using the `curve_fit` routine from the `scipy.optimize` module, a standard implementation of the least-sum-of-squares fit routine. To extract characteristic timescales of fluorescence recovery and decay, as well as associated stable fractions, FRAP and FDAP curves were respectively fitted with the following functions: For normalised FRAP intensity, we used 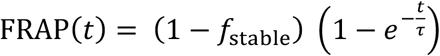 with characteristic time constant *τ* and stable fraction *f*_stable_, whereas for normalised FDAP intensity, we introduced an additional offset *f*_off_ to account for the observed overactivation in the first time frame after photoconversion, and used 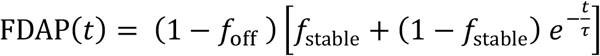.

## Results

### Exchange kinetics of GlyRs at synapses in spinal cord neurons

An experimental FRAP/FDAP protocol was established to simultaneously quantify the recruitment and the loss of GlyRs and gephyrin at inhibitory synapses (see Methods, Fig. S2, S3). Receptors and scaffold proteins were tagged with the photoconvertible fluorophore Dendra2 and expressed in cultured spinal cord neurons using lentivirus infection. Dendra2 was photoconverted with a 405 nm laser focussed on a single synaptic punctum, followed by time-lapse imaging to record the fluorescence recovery after photobleaching (FRAP) of the green (non-converted) fluorescence over 30 minutes. Concurrently, we measured the fluorescence decay after photoconversion (FDAP) in the red channel (photoconverted Dendra2) as an additional read-out of the protein dynamics.

In living neurons, the GlyR associated fluorescence of the bleached puncta recovered from close to background levels at t = 0 to about 60% of its baseline value after 30 minutes (Fig. 1). We noticed that the Dendra2-GlyRα1 signals in the area surrounding the bleached punctum increased after the 405 nm pulse due to the low intensity halo of the laser. The increase was much less pronounced at remote areas (far controls). To compensate this photochromic effect of Dendra2, the intensity data were normalised using control puncta in the proximity of the bleached spot (near controls, see Methods for FRAP data analysis).

**Figure 1.**
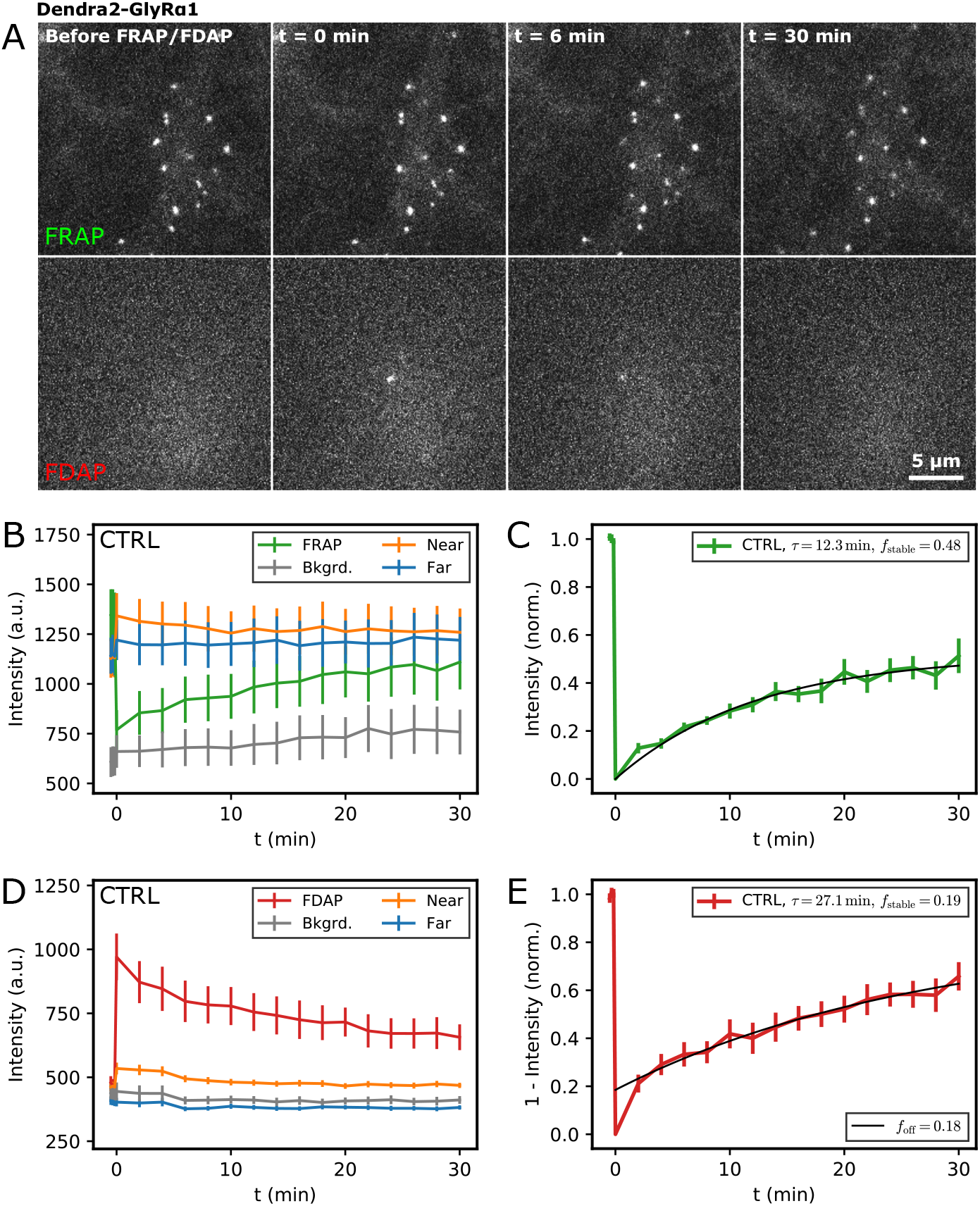
Dendra2-GlyR dynamics in living neurons. (A) FRAP/FDAP recordings of Dendra2-GlyRα1 in living spinal cord neurons. Representative time-lapse images from left to right: 1. before photoconversion, 2. t = 0 (after exposure to a focussed 405 nm laser beam), 3. t = 6 min, 4. t = 30 min. Top: FRAP of nonconverted (green) Dendra2-GlyRα1, bottom row: FDAP of photoconverted (red) Dendra2-GlyRα1. (B) FRAP quantification of the average fluorescence intensity (a.u.) of synaptic puncta of non-converted Dendra2-GlyRα1 (mean ± SEM, n = 14). For analysis, the recovery of the exposed puncta (FRAP, green trace) was normalised by the fluorescence intensity of neighbouring control puncta (near controls, orange) that exhibit a slight photochromic activation, not visible in control puncta at the edge of the field of image (far controls, blue). The grey trace represents the average background intensity of diffuse Dendra2-GlyRα1 in the extrasynaptic plasma membrane. (C) FRAP dynamics of Dendra2-GlyRα1 (normalised data) were fitted with a single exponential component τ and a stable fraction *f*stable. (D) FDAP average fluorescence intensity (a.u.) of synaptic puncta of photoconverted Dendra2-GlyRα1 (mean ± SEM, n = 14). For FDAP analysis (E), the recovery of the photoconverted puncta (FDAP, red trace) was normalised by the fluorescence intensity prior to phoconversion (set to zero) and after photoconversion (set to 1).

As expected, the red fluorescence of photoconverted Dendra2-GlyRα1 at synaptic puncta decreased in parallel to the recovery of the green fluorescence (Fig. 1). The loss of fluorescence was similar to the rate of recovery, falling to about 34% of the initial value after 30 minutes. Control puncta that were near the site of photoconversion also showed a slight increase in fluorescence in response to the 405 nm laser pulse, confirming that stray light can affect the fluorophores despite its low intensity. FDAP intensity traces were normalised and rescaled as described in the Methods section.

Our pilot experiments had shown that the recovery and the loss of Dendra2-gephyrin was roughly on the order of 50% after 30 minutes (Fig. S3). In other words, GlyR and gephyrin populations exchange on a similar timescale, which we thought could be an indication that the two components enter and exit synapses jointly in the form of GlyR-gephyrin complexes, consistent with earlier hypotheses (10, 19). We reasoned that if this was true, the immobilisation of the GlyRs should reduce the exchange rate of gephyrin at synapses.

### GlyR immuno-immobilisation (IMMO)

To interfere with the mobility of the GlyRs, we decided to crosslink the cell surface receptors using specific antibodies against the α1-subunit of the GlyR. Antibody crosslinking has been previously shown to fully block the lateral diffusion of neurotransmitter receptors at excitatory synapses (23). Using the same approach, spinal cord neuron cultures were treated for 10 minutes with high concentrations of primary antibodies against GlyRα1, followed by a 10 minute application of Alexa Fluor 488 (A488) conjugated secondary antibodies. Since the Dendra2 fluorophore can mask the GlyRα1 epitope (supplementary information in (7)), these experiments were performed on naive neurons expressing endogenous GlyRs. FRAP was then carried out on the A488 dyes attached to the crosslinked endogenous GlyRs using a 488 nm laser (Fig. 2).

**Figure 2.**
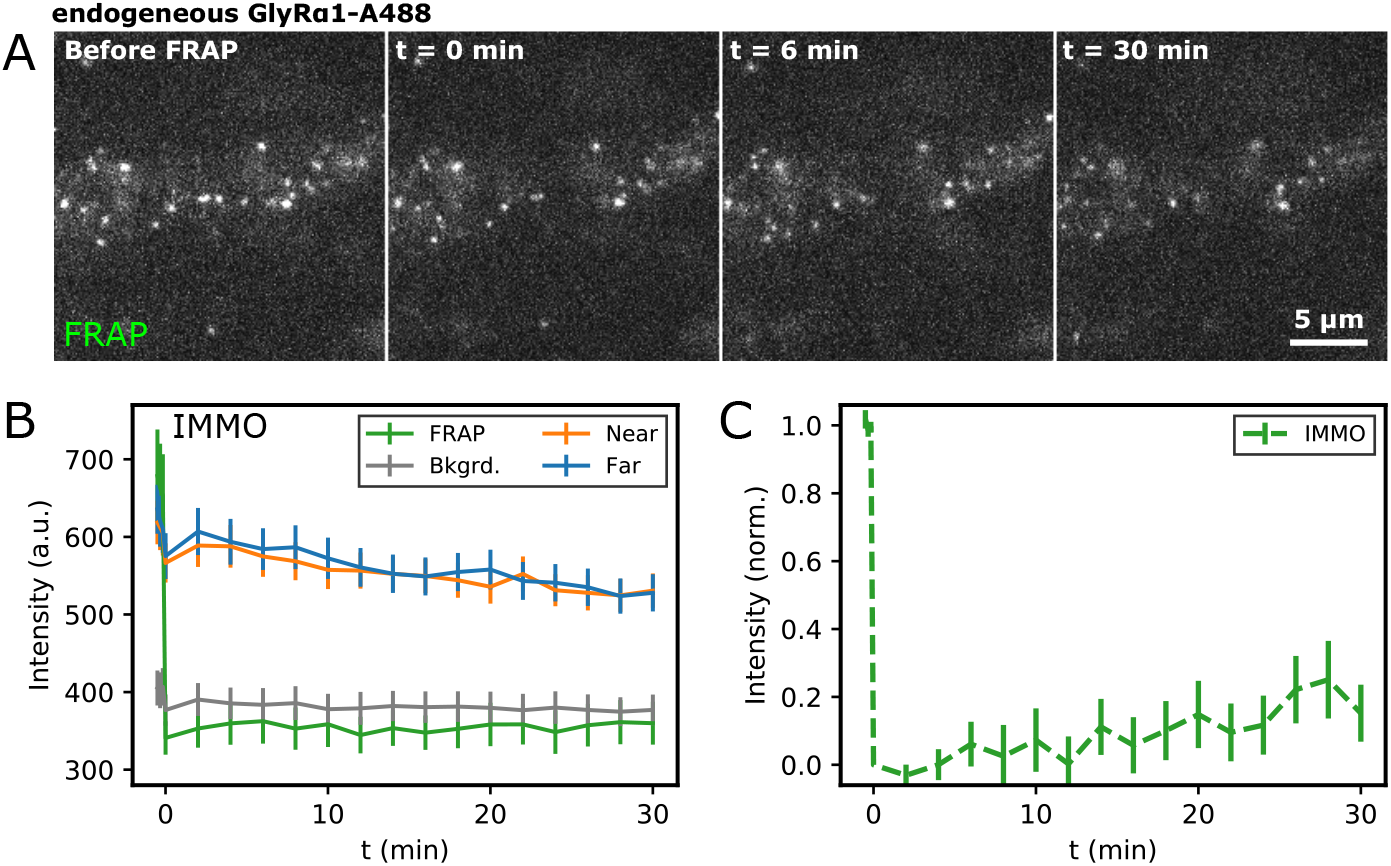
The effect of immuno-immobilisation on GlyR dynamics. (A) FRAP of immuno-immobilised GlyRs in living spinal cord neurons using a focussed 488 nm laser pulse. Endogenous GlyRs were incubated sequentially with rabbit anti-GlyRα1 and anti-rabbit secondary antibodies conjugated with A488 dye. (B) FRAP data analysis and quantification of the exchange dynamics of immuno-immobilised endogenous GlyRs (mean ± SEM, n = 13). (C) Normalised FRAP data.

Antibody binding blocked the fluorescence recovery at GlyR puncta almost entirely. In absolute terms, the fluorescence intensity after bleaching of the A488 dyes remained at background levels throughout the recordings (Fig. 2). After normalisation of the data, a minor recovery could be discerned, however, this is likely the result of the more pronounced photobleaching of the A488 fluorophores during image acquisition (control puncta). Nonetheless, it can be concluded that crosslinking had a dramatic effect on the mobility of endogenous GlyRs when compared to the exchange rates of recombinant Dendra2-GlyRα1 containing complexes under control conditions (Fig. 1). We did not observe any obvious differences in the sub-cellular distribution of GlyRs after crosslinking, as judged by immunocytochemistry using the vesicular inhibitory amino acid transporter (VIAAT) as presynaptic marker (Fig. S4). Both endogenous GlyRs and Dendra2-gephyrin showed extensive co-localisation with VIAAT in the control condition as well as after immuno-immobilisation.

### Effects of GlyR immuno-immobilisation on the exchange kinetics of gephyrin

Having demonstrated the efficacy of GlyR crosslinking, we examined the consequences of receptor immuno-immobilisation (IMMO) on the dynamics of the synaptic gephyrin scaffold. To do so, we performed FRAP/FDAP experiments in spinal cord neurons expressing Dendra2-gephyrin. Endogenous GlyRs were immuno-immobilised as before using primary GlyRα1 antibody and A647-conjugated secondary antibody. Control neurons were treated in the same way in these experiments, only that the antibodies were omitted during the incubation in Tyrode solution (Fig. 3, S5).

**Figure 3.**
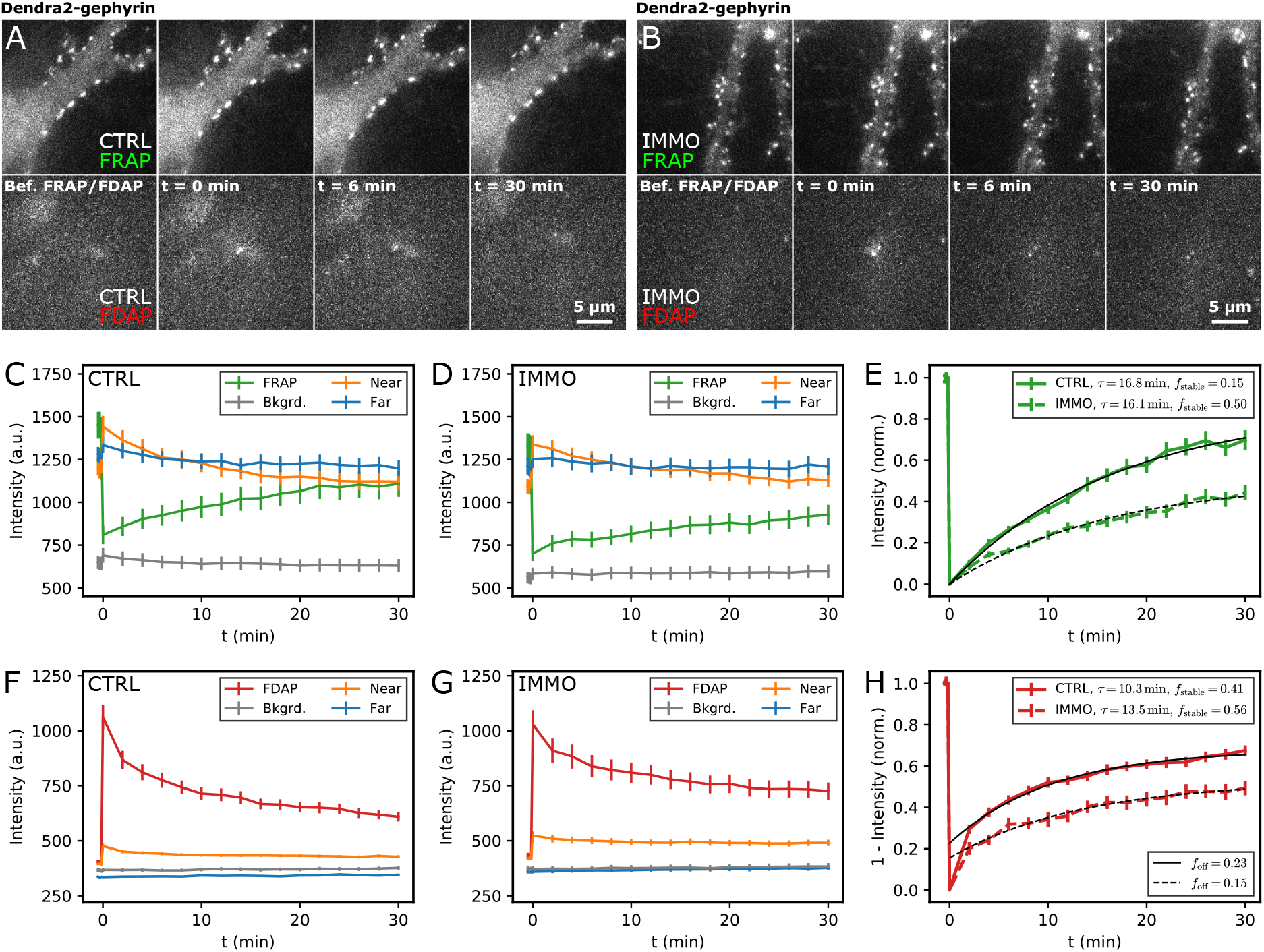
Dendra2-gephyrin dynamics in control and immuno-immobilised conditions. (A,B) FRAP/FDAP time-lapse images of Dendra2-gephyrin in spinal cord neurons under control conditions (A) and after GlyR immuno-immobilisation (B). Top: FRAP of nonconverted (green) Dendra2-gephyrin, bottom row: FDAP of photoconverted (red) Dendra2-gephyrin. (C,D) Quantification of the average intensity of Dendra2-gephyrin at synaptic puncta in FRAP recordings in control (C, mean ± SEM, n_CTRL_ = 48) and immuno-immobilised conditions (D, mean ± SEM, n_IMMO_ = 55). (E) Normalised FRAP data fitted with a single exponential component τ and a stable fraction *f*_stable_. (F,G) Quantification of FDAP recordings of Dendra2-gephyrin (F, n_CTRL_ = 48; G, n_IMMO_ = 55, mean ± SEM). (H) Normalised and fitted FDAP data.

The recovery of the green Dendra2-gephyrin fluorescence was substantially reduced after crosslinking of the receptors compared to the control condition (44% vs 70% of the baseline after 30 min; Fig. 3). The normalised FRAP curves recorded over a period of 30 minutes were fitted with two free parameters, a time constant τ and a weighing factor *f* that describes the fraction of fluorophores in the stable pool. Interestingly, fitting of the FRAP data showed that the time constant of the recovery was not significantly different between the two conditions (τ_CTRL_ = 16.8 ± 1.2 min, τ_IMMO_ = 16.1 ± 1.8 min; 95% confidence interval reported from fit routine), but that the stable fraction of Dendra2-gephyrin was increased by GlyR crosslinking from 0.15 to about 0.5. These observations were confirmed by the decay of the red fluorescence (FDAP). Again, immuno-immobilisation increased the fraction of stable fluorescence (*f*_CTRL_ = 0.41 ± 0.01, *f*_IMMO_ = 0.56 ± 0.02), but had only a minor effect on the decay rate (τ_CTRL_ = 10.3 ± 0.9 min, τ_IMMO_ = 13.5 ± 2.0 min). It should be noted that the intensity in the red channel at the first time point after photoconversion was systematically higher than in the subsequent images, which is why we fitted an additional offset correcting for the overactivation at t = 0 (see Methods).

To exclude the existence of a faster component that was not captured with a 2 min acquisition frequency, we conducted another series of FRAP/FDAP experiments with a higher acquisition rate (15 s over a period of 4 min). The Dendra2-gephyrin intensity showed only a small recovery and decay on this timescale. Moreover, the data could be approximated with a linear fit, indicating that no sizeable fast component of exchange was present (Fig. S5). However, there was a trend that the gephyrin exchange was reduced by GlyR immuno-immobilisation (recovery slope *a*_CTRL_ = 0.067 ± 0.004 min^-1^, *a*_IMMO_ = 0.055 ± 0.003 min^-1^).

The fit of the FRAP/FDAP data recorded over a period of 30 minutes with two parameters (τ, *f*) implies that there is a seemingly immobile fraction of gephyrin that does not exchange with the extrasynaptic pool on this timescale. The fact that immuno-immobilisation increases this immobile fraction suggests that the stable gephyrin population is dependent on receptor-scaffold interactions at synapses. In other words, it is possible that immobile GlyRs provide stable high-affinity binding sites for gephyrin at inhibitory synapses.

### Effects of GlyR immuno-immobilisation on synaptic receptor and gephyrin levels

Given that the size of synaptic gephyrin domains is thought to depend on the balance between GlyR-mediated diffusion and capture of extrasynaptic GlyR-gephyrin complexes and the loss of synaptic gephyrin into the cytoplasm by desorption (19), we asked whether GlyR and gephyrin levels would be affected by immuno-immobilisation of the receptor. We therefore compared the intensities of synaptic puncta at the beginning (before FRAP/FDAP) and at the end of our recordings (t = 30 min; Fig. 4). We first verified that immuno-immobilisation did not change synaptic GlyR levels, indicating that a stationary state was reached at the end of the treatment protocol. Indeed, the intensity of crosslinked GlyR puncta remained stable throughout the recording (I_pre_ = 1206 ± 911 and I_30min_ = 1131 ± 961 a.u., mean ± SD, n = 336 from 54 cells, MW test p = 0.114; Fig. 4B). We then measured the average intensities of all clearly identifiable gephyrin puncta in the CTRL and IMMO conditions at both time points. The average intensity of gephyrin puncta was not different between the two conditions, suggesting that synaptic size is not dependent on GlyR-mediated gephyrin dynamics on the timescale of the experiment. The average intensity of synaptic puncta at the start of the FRAP/FDAP acquisition (approx. 5-15 minutes after immuno-immobilisation) was I_CTRL_ = 1279 ± 825 and I_IMMO_ = 1240 ± 737 a.u. (n_CTRL_ = 3031, n_IMMO_ = 3318, MW p = 0.034). At the end of the recordings (approx. 40-55 min after treatment) the average intensities were I_CTRL_ = 1322 ± 933 and I_IMMO_ = 1331 ± 1059 a.u. (n_CTRL_ = 3261, n_IMMO_ = 3640, MW p = 0.48).

**Figure 4.**
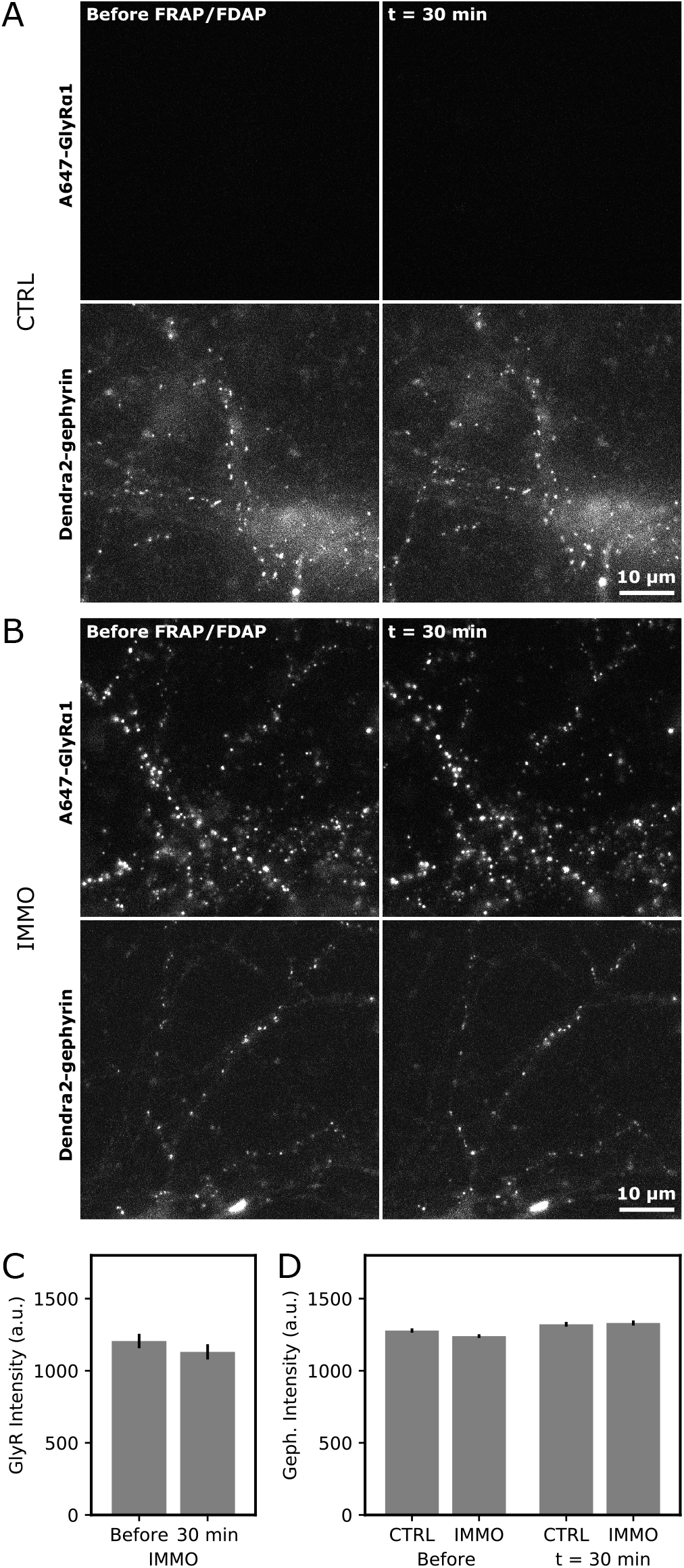
Cluster intensities after GlyR immuno-immobilisation. (A) Time-lapse images of Dendra2-gephyrin in the control condition taken at the beginning (before photoconversion) and the end of the 30 min FRAP recordings (lower panels, green channel). The upper panels show the empty far red channel (control condition without GlyR crosslinking). (B) Time-lapse images of immuno-immobilised GlyRs (top panels, far red channel) and Dendra2-gephyrin (bottom, green channel) taken at the beginning (before FRAP/FDAP) and at the end of the recording (30 min). (C) Quantification of the average fluorescence intensity of A647-GlyRα1 at identified synaptic puncta at the beginning and at the end of the FRAP recordings (IMMO condition, nbefore = 336, n30min = 336 from 54 cells). (D) Quantification of all Dendra2-gephyrin puncta in control and immuno-immobilised conditions at the beginning and at the end of the FRAP recording (n_CTRL_,before = 3031, n_CTRL_,30min = 3261 from 56 fields of view; n_IMMO_, before = 3318, n_IMMO_, 30min = 3640 from 57 fields of view).

### Model of receptor and scaffold dynamics at synaptic complexes

To integrate the different experimental data and to gain further insight into the interdependent kinetics of synaptic GlyRs and gephyrin, we devised a simple model of receptor-scaffold dynamics at inhibitory synapses (Fig. 5, SI Text). For simplicity, we consider only three species or molecular states at the synapse: Receptors *r* that are diffusing and/or transiently attached to loosely interacting scaffold proteins *s*, and a population of more tightly bound receptor-scaffold complexes *c*.

**Figure 5.**
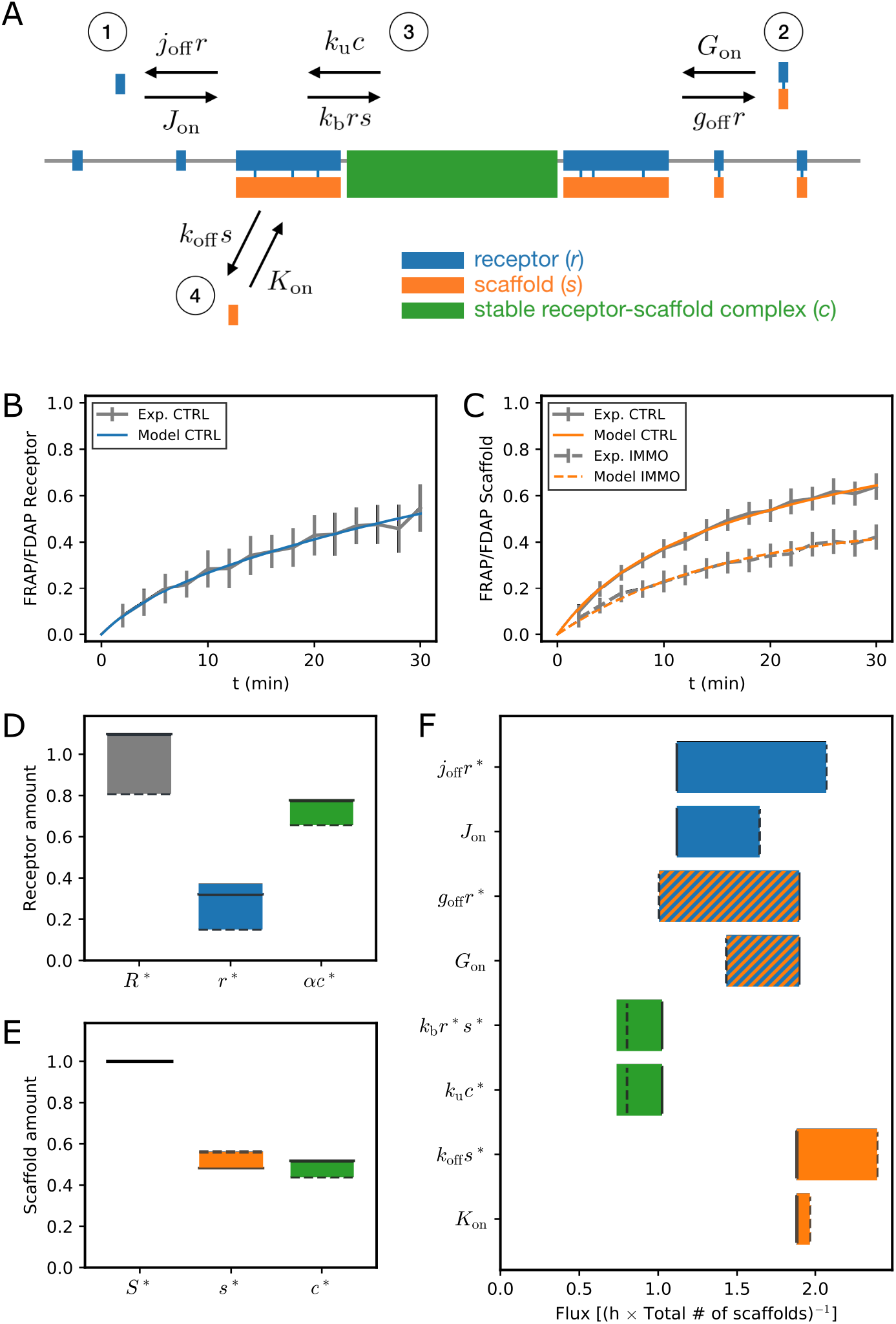
Model of synaptic receptor and scaffold protein dynamics. (A) Sketch of the proposed exchange kinetics. Receptors can enter and exit the synapse alone (1) or with scaffold proteins attached (2). Synaptic receptors and scaffold proteins exist either as loosely bound populations (receptors *r*, blue, and scaffold proteins *s*, orange) or in a more tightly bound state (‘crosslinked’ receptor-scaffold complexes *c*, green); the transitions between the respective states are described by the fluxes (3). Scaffold proteins exchange with the cytoplasm (4) when not in the tightly bound state. (B,C) Fit of the combined experimental FRAP/FDAP curves (see Methods and SI Text) for the GlyR dynamics in the control condition (B) and gephyrin dynamics in the control and immuno-immobilised conditions (C). The model curves correspond to the best fit parameters for *f* = 0. (D,E) Total amount and breakdown into subpopulations of synaptic receptors (D) and scaffold proteins (E) predicted by the model, relative to the total number of scaffold proteins at the synapse (S* = 1). The bars indicate the range of values obtained for 0 ≤ *f* ≤ 1; values for *f* = 0 and *f* = 1 are indicated by solid and dashed lines, respectively. Note that in the model one receptor represents a pentameric GlyR complex, and one scaffold particle represents one gephyrin trimer. The respective contributions of the loosely and tightly bound populations are color-coded according to panel (A). (F) Receptor and scaffold fluxes predicted by the model in the control condition. The bars indicate the range of values obtained for 0 ≤ *f* ≤ 1; values for *f* = 0 and *f* = 1 are indicated by solid and dashed lines, respectively. Note that in the immuno-immobilised condition all receptor fluxes are assumed to vanish, and *k*_off_*s* = *K*_on_.

The dynamics of receptor and scaffold populations arise from transitions between the states *r, s*, and *c*, and incoming as well as outgoing protein fluxes (Fig. 5A). Loosely bound receptors *r* and scaffolds *s* exchange with extrasynaptic pools, where receptors enter the synapse with a flux *J*_on_ and exit with a rate *j*_off_ into the extrasynaptic membrane. Scaffolds are recruited from and exit into the cytoplasm with an influx *K*_on_ and at a rate *k*_off_, respectively. In addition, extrasynaptic receptor-scaffold complexes enter into the synapse with a flux *G*_on_, and individual receptor-scaffold complexes are released into the extrasynaptic membrane with a rate *g*_off_. Inside the synapse, loosely bound receptors and scaffolds can form a more tightly crosslinked state *c* with an effective binding rate *k*_b_; inversely, more stable receptor-scaffold complexes in state *c* can give way to loosely bound receptors *r* and scaffolds *s* with an effective unbinding rate *k*_u_. For this reaction, we assume a fixed stoichiometry *α* = 1.5 between receptors and scaffolds in state *c* based on GlyR and gephyrin trimer properties, see SI Text. In principle, the model is then completely characterized by the 8 parameters *J*_on_, *j*_off_, *K*_on_, *k*_off_, *G*_on_, *g*_off_, *k*_b_, and *k*_u_. At the stationary state, receptor and scaffold in- and effluxes balance each other, which allows to determine the stationary values of all considered populations as a function of the model parameters (SI Text). By normalising receptor and scaffold amounts by the total amount of scaffolds at the synapse, we can express *K*_on_ as a function of all other parameters.

We mimicked the immuno-immobilisation protocol by setting all fluxes involving receptors to zero, as they cannot enter nor exit from the synapse in the immobilised condition. Since we cannot exclude that antibody-mediated crosslinking of diffusing or loosely bound receptors in state *r* affects their synaptic organization, we furthermore admit that a fraction *f* of these receptors eventually end up in the highly interacting receptor-scaffold complex *c*. We present here the results covering the complete range 0 ≤ *f* ≤ 1. Based on the experimental observation that the amount of scaffold proteins does not change among conditions, we require that the total amount of scaffolds *S = s + c* is constant, which imposes an additional constraint on the parameters and reduces the number of free parameters to six (see SI Text).

Our model then allows to predict the time course of FRAP/FDAP experiments for receptors in the CTRL and scaffold proteins in the CTRL and IMMO conditions as a function of the model parameters (see SI Text). In order to determine the parameters of the model, we tried and fitted the predicted time courses to the experimentally obtained time courses (Fig. 5B,C for *f* = 0). Based on these fits, we can quantify the total amount of synaptic receptors (*R*) as well as the different contributions of loosely bound (*r, s*) or more strongly interacting (*c*) receptor and scaffold protein populations (Fig. 5D,E and Fig. II in SI Text). Our model predicts the ratio of receptors to scaffold proteins to be 0.8-1.1 (Fig. 5D), which corresponds to a ratio of 0.3-0.4 pentameric GlyRs per gephyrin monomer, since we take gephyrin trimers to be the basic unit of scaffold proteins in the model. Furthermore, the model suggests that in the control condition, only a small fraction (~30%) of receptors is loosely bound and exchanges with the extrasynaptic membrane, while the majority of receptors exists in the stable state (Fig. 5D). According to the model, scaffold proteins are more equally distributed between the loosely bound and the stable states (Fig. 5E).

We can furthermore ask what are the respective fluxes of receptors and scaffolds exiting and entering the synapse, as well as the fluxes related to binding and unbinding of receptors *r* and scaffolds *s* into the more stable complex *c* (Fig. 5F). Our model suggests that the lateral influx *G*_on_ of receptor-scaffold complexes is similar to the influx *J*_on_ of receptors that enter the synapse without a scaffold protein attached, and the same holds for the exiting fluxes *j*_off_ *r*\* and *g*_off_ *r*\*, respectively. The cytoplasmic recruitment *K*_on_ of scaffold proteins tends to outweigh the lateral receptor-mediated scaffold influx *G*_on_; the loss *k*_off_ *s*\* of scaffolds to the cytoplasm is equal to *K*_on_ if not somewhat larger. The exchange between the loosely bound states *r* and *s* and the highly interconnected state *c* is considerably slower than all receptor and scaffold exchanges with extrasynaptic pools.

While for *f* = 0 all fluxes are individually balanced (i.e., *J*_on_ = *j*_off_ *r*\*, *G*_on_ = *g*_off_ *r*\* etc., see SI Text), we cannot exclude that there is a net influx of scaffold proteins arriving in the form of GlyR-gephyrin complexes by lateral membrane diffusion, as we find *G*_on_ > *g*_off_ *r*\* for *f* > 0 (Fig. 5F, Fig. III in SI Text). A net influx would violate detailed balance and thus imply a departure from thermodynamic equilibrium; the synapse would be in a non-equilibrium stationary state. Our model shows, however, that the cytoplasmic recruitment of scaffold proteins at the synapse contributes in all cases significantly to the renewal of synaptic scaffolds in the control condition, and any departure from thermodynamic equilibrium would supposedly be minor.

## Discussion

Our data disclose the reciprocity of receptor and scaffold protein dynamics and clustering at glycinergic spinal cord synapses, mediated by strong interactions between the GlyR β-subunit and the synaptic scaffold protein gephyrin. A biophysical model of our data identified different degrees of receptor stability at synapses; on the one hand a more loosely interacting population of receptors and scaffold proteins, and on the other hand a more a tightly complexed receptor-scaffold network.

### FRAP/FDAP of gephyrin and GlyRs

To determine the dynamic behaviour of gephyrin and GlyRs in cultured spinal cord neurons, we established an analytical protocol based on the photoconversion and time-lapse imaging of fluorescently tagged recombinant proteins at synapses. The photoconvertible fluorophore Dendra2 was chosen for our experiments, since it displays less photochromism in the green channel in response to near UV illumination as opposed to mEos4b (24). Local reference puncta (near control points) were used to correct photochromic effects and the photobleaching during image acquisition.

There is some evidence that the overexpression of recombinant GlyRs and gephyrin does not substantially alter the copy numbers at synapses ((7, 25), supplementary data). In the case of the receptor, synaptic targeting is strictly dependent on the assembly of Dendra2-GlyRα1 with the endogenous β-subunit. This suggests that the overexpression of recombinant GlyRα1 replaces the majority of the endogenous α-subunits without changing the absolute numbers at synapses. Even though it cannot be entirely ruled out that the overexpression and fluorescent tagging may have some impact on the synaptic structure, the synapses have most likely reached a steady state by the time the FRAP/FDAP experiments are carried out, given that lentiviral infection was done several days prior to synaptogenesis.

Our experimental results are largely consistent with earlier studies of the population dynamics of gephyrin and GlyRs. For instance, synaptic puncta of transfected Venus-gephyrin and mRFP-gephyrin, as well as endogenous (knock-in) mRFP-gephyrin in cultured spinal cord neurons recover to about 40% of their initial fluorescence within 30 minutes (15). A relatively broad distribution of time constants on the order of hours was determined for synaptic Dendra2-GlyRα1 in motoneurons of transgenic zebrafish larvae (26). It is noteworthy that the FRAP/FDAP traces in our experiments were fitted with a single exponential recovery/decay, alongside a much slower component that we considered as stable within the duration of our recordings. It is therefore expected that the characteristic timescales obtained from our fits are faster than the respective timescales of a complete recovery.

In organotypic hippocampal slices, the rate of recovery of endogenous EGFP-gephyrin clusters was shown to be highly variable (27). In addition to a subtle size-dependence of the exchange rate, the authors observed a strong developmental stabilisation of gephyrin. However, these data are not directly comparable to our situation, since inhibitory synapses in the hippocampus are overwhelmingly GABAergic and likely follow different clustering mechanisms (1).

### The effect of receptor immuno-immobilisation on scaffold protein dynamics

On a purely qualitative level, our data reveal that GlyR crosslinking reduces the exchange of Dendra2-gephyrin at synapses. A possible explanation could be that the dynamics of gephyrin depends to a certain extent on the entry and exit of GlyR-gephyrin complexes at inhibitory synapses. This is due to the fact that the GlyRβ-gephyrin interaction is remarkably stable, allowing extrasynaptic GlyR-gephyrin complexes to integrate into the synaptic scaffold via multiple interaction sites (GlyR-gephyrin and gephyrin-gephyrin) (1). At excitatory synapses, immuno-immobilisation of AMPA receptors did not produce a slowdown of the exchange rate of the scaffold protein βSAP97 (23), which is consistent with a role of βSAP97 in the forward trafficking of AMPA receptors to the plasma membrane, but not their integration into the synaptic membrane (28).

Another possible explanation would link the reduction of gephyrin exchange to the crosslinking and immobilisation of the synaptic GlyR population. In this scenario, immobile GlyRs form stable interactions with synaptic gephyrin molecules that are thus prevented from exchanging with extrasynaptic pools. In line with this interpretation, GlyR crosslinking did not change the steady state level of gephyrin at synapses, pointing to a mutual stabilisation between receptors and scaffold proteins at the synapse.

### A model of reciprocal GlyR-gephyrin stabilisation

Based on our experimental observations, we aimed to develop a biophysical model that would provide quantitative insight into the synaptic organisation and dynamics of GlyRs and gephyrin molecules beyond the apparent stable fractions and characteristic timescales. We propose a model in which the extrasynaptic receptor and scaffold pools exchange with loosely bound populations at synapses, that are in turn in a dynamic equilibrium with a tightly interacting receptor-scaffold complex. This simple model is sufficient to account for all experimental FRAP/FDAP curves, where the crosslinking of GlyRs is mimicked by a suppression of all GlyR-associated fluxes.

Interestingly, our model does not require the existence of a fully stable component of synaptic GlyR or gephyrin, as one could infer from the heuristic fits of a single exponential decay to our experimental data. The model instead suggests that the observed dynamics arise from the interplay between the fast exchange of loosely bound synaptic populations and extrasynaptic pools on the one hand, and their slow exchange with the tightly interacting synaptic receptor-scaffold complex on the other hand. The existence of a synaptic component with a higher degree of stabilisation had been previously proposed based on a large fraction of GlyRs that do not appear to swap between synaptic and extrasynaptic locations on a timescale of minutes (10). To what extent the stable synaptic population relies solely on receptor-scaffold interactions or depends on additional factors such as gephyrin palmitoylation or binding to adhesion proteins cannot be decided at this stage (1).

While in the model all receptor and scaffold quantities are expressed in terms of the total amount of scaffold proteins, we can obtain an estimate of the copy numbers of synaptic GlyRs from the typical number of synaptic gephyrin trimers. If we assume the latter to be ~100 (corresponding to ~300 gephyrin monomers/synapse on average), the model predicts the number of synaptic GlyRs to be of the order of 80-110, closely matching earlier experimental results (7). A majority of these (~70%) interact tightly with synaptic gephyrin, whereas the remainder is more loosely bound and exchanges with extrasynaptic pools. We can furthermore obtain estimates of the number of extrasynaptic GlyRs and gephyrins that enter the synapse per unit of time: For GlyRs, the flux amounts to ~5 GlyRs/minute, and we obtain a slightly larger value for gephyrin with ~6 gephyrin trimers/minute.

### Single molecule dynamics predicted by our model

Although our model relies on a coarse-grained description of receptor-scaffold dynamics at synapses, it allows predictions about single molecule dynamics that go beyond the scope of this study. The existence of a small receptor population that exchanges with the extrasynaptic pool on a fast timescale implies that individual receptor dwell times can be considerably shorter than the typical time constant of GlyRs observed in our FRAP experiments (Fig. 6). Our model predicts that a considerable fraction (10-30%) of GlyRs leave the synapse in less than a minute, as opposed to a fluorescence recovery of comparable size in up to ten minutes. The prediction of short receptor dwell times is consistent with the literature, where on very short timescales (< 40 s) a distinction between swapping (exchanging) and stable receptors has been made, and where typical dwell times for the swapping population have been determined (e.g. (29)). However, a more detailed quantitative comparison with single molecule data is hampered by experimental limitations such as the size of quantum dots (30) or the insufficient localisation precision of single molecule diffusion data (31).

**Figure 6.**
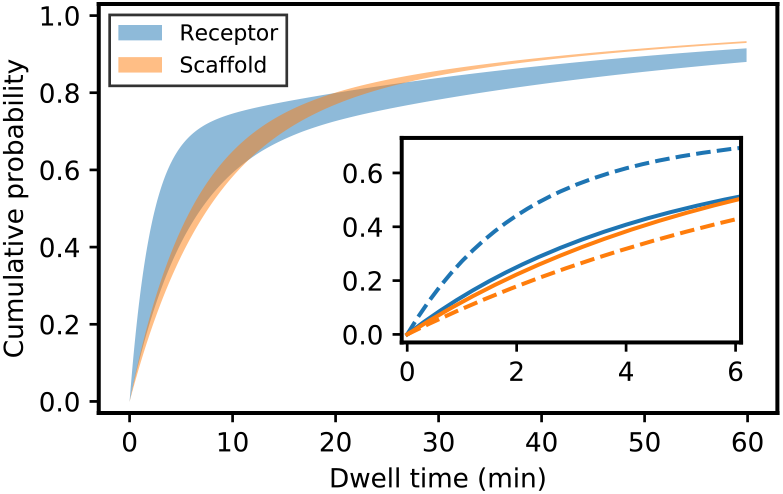
Predicted dwell times of receptors and scaffolds after entry into the synapse. While the characteristic FRAP/FDAP timescales of both receptors (blue) and scaffold proteins (orange) are similar and of the order of tens of minutes (Fig. 1, 3), a sizeable proportion of receptors have dwell times below a minute, as opposed to those of scaffold proteins (inset). Shaded bands indicate the range of values obtained for 0 ≤ *f* ≤ 1; solid and dashed lines represent the predicted dwell times for *f* = 0 and *f* = 1, respectively.

### Relation to previous models of receptor-scaffold organisation at inhibitory synapses

Earlier modelling studies of receptor-scaffold organisation at inhibitory synapses did not explicitly address the exchange kinetics of synaptic GlyRs and gephyrin, but we can try to assess the consequences of GlyR immobilisation for each of them. In a recent study, we hypothesised that the size of the postsynaptic domain is maintained by the recruitment of extrasynaptic GlyR-gephyrin complexes (here denoted by *G*_on_) that replace gephyrin molecules that are lost due to desorption (*k*_off_ *s*\*) (19). In our model, we considered that these fluxes dominated the lateral outward flux of scaffolding proteins bound to receptors (*g*_off_ *r*\*) as well as the incoming flux from the cytoplasm (*K*on). In this limit, the resulting nonequilibrium stationary state depends on the diffusion of GlyR-gephyrin complexes in the extrasynaptic membrane; the immobilisation of GlyRs should therefore lead to the depletion of synaptic gephyrin. This size decrease is not seen on the timescale of the present experiments (Fig. 4). The incoming lateral flux *G*_on_ and the outgoing flux *k*_off_ *s*\* determined in the present work are comparable to what we estimated earlier (19). However, the receptor-mediated lateral efflux *g*_off_ *r*\* is found to be comparable to the corresponding influx *G*on, and similarly the scaffold influx from the cytoplasm (*K*_on_) is found to compensate the losses due to scaffold desorption into the cytoplasm (*k*_off_ *s*\*), as would be expected if the postsynaptic domain was a structure at or close to thermodynamic equilibrium.

Our data therefore strongly suggest that the size of the synaptic gephyrin domain is determined by processes other than the simple recruitment of GlyR-gephyrin complexes, although it cannot be ruled out that more GlyR-gephyrin complexes enter the synapse (*G*_on_ in the model) than leave the synapse (*g*_off_ *r*\*) (Fig. 5). However, our findings support the conclusion that all lateral fluxes contribute to the synaptic dynamics, since a model in which *G*_on_ and *g*_off_ *r* are not considered does not fit the experimental data satisfactorily (SI Text Appendix B). It remains to be mechanistically understood how the interaction between receptors and scaffold proteins with different degrees of stabilisation, as well as their interactions with other synaptic proteins, contribute to synaptic size regulation.

It appears less straightforward to interpret our results in light of the reaction-diffusion model proposed by Haselwandter and colleagues (17, 21). The authors identified several key reactions necessary for the spontaneous formation of scaffold domains, notably the recruitment of cytoplasmic scaffold proteins and cytoplasmic receptors (exocytosis) by synaptic scaffold proteins. These reactions are limited by steric repulsion of proteins, and incoming cytoplasmic fluxes are balanced by diffusive fluxes of receptors and scaffolds into the extrasynaptic membrane. The effect of crosslinking of GlyRs crucially depends on how the various reactions are modified in this setting, and cannot be *a priori* estimated from the model equations. Because the size of the domains is entirely dependent on the interplay of receptor and scaffold spatio-temporal dynamics, however, receptor immobilisation should significantly affect scaffold domain size in this model, which is not supported by our data. The Turing instability proposed to be at the root of postsynaptic domain size determination is intrinsically a non-equilibrium phenomenon and appears also at odds with the (at least approximate) balance of individual fluxes, a hallmark of thermodynamic equilibrium.

In contrast, our experimental results and the proposed model are broadly consistent with the quasi-equilibrium model of Sekimoto and Triller (18), who predicted that condensed phases of scaffolds and receptors would arise spontaneously when the interaction between the two is sufficiently strong (see also (32)). In this model, the size of the domains is externally controlled, and the nucleation of the condensed phase constrained to the synaptic area by additional molecular interactions at the synapse, in line with the kinetic model presented here. While this earlier study convincingly argued that the reciprocal stabilisation of scaffolds and receptors may support the formation of stable postsynaptic domains, we present here a detailed, quantitative account of the underlying reaction kinetics, fluxes, and synaptic organisation.

Other models of synaptic scaffold protein dynamics have been proposed that did not specifically address inhibitory synapses or the role of receptors. Shomar *et al*. (33) argued that both scaffold recruitment from and desorption of scaffolds into the cytoplasm are cooperative processes, which would allow to explain the observed skewed distributions of synapse sizes (34). However, this model cannot account for the observed changes in the FRAP/FDAP traces after receptor immobilization, and it is not clear from our data that scaffold binding and unbinding have to be cooperative processes.

### Conclusion and perspectives

The quantitative analysis of excitatory and inhibitory synaptic size dynamics has received much attention in recent years (reviewed in (35)), and concomitant theoretical modelling ranged from very generic statistical (34) to more biophysical models of synaptic scaffold dynamics (19, 33). However, an integrated account of synaptic size dynamics that takes into account both receptors and scaffold proteins has so far been lacking. While the large number of different molecular players at synapses (e.g. (36-38)) precludes a microscopically detailed biophysical model involving all relevant species in the foreseeable future, our model with three distinct receptor and scaffold populations is a first step towards a more comprehensive picture of glycinergic synapse dynamics. Although we did not explicitly address synaptic size fluctuations in this work as we restricted our analysis to the average dynamics with a stationary synaptic size, it would be straightforward to extend our model to account for molecular fluctuations.

In this work, we identified loosely bound glycine receptor and gephyrin scaffold populations that co-exist with a stable receptor-scaffold complex at inhibitory synapses. It is tempting to speculate that these mobile and stable populations differentially contribute to the plasticity and the stability of glycinergic synapses. More generally, it will be interesting to explore to what extent similar descriptions may apply to excitatory synapses that are also stabilized by scaffold proteins interacting with mobile receptors.

## Acknowledgements

Research in our laboratory is funded by the European Research Council (ERC, Plastinhib), Agence Nationale de la Recherche (ANR, Synaptune and Syntrack), Labex (Memolife) and France-BioImaging (FBI). We are grateful to Xiaojuan Yang and Astou Tangara for technical help and Noam Ziv (Technion, Israel Institute of Technology) for the Openview software.

## Author contributions

VH, AT, JR and CGS designed the research; TC and CGS planned and performed the experiments; TC, JR and CGS analysed the data; JR developed and analysed the model with input from all authors; TC, JR and CGS wrote the manuscript; all authors reviewed and edited the final manuscript.

