## Supplementary Methods and Figures for "Reciprocal stabilisation of glycine receptors and gephyrin scaffold proteins at inhibitory synapses"

#### Expression constructs and lentivirus preparation

Full-length gephyrin tagged with various fluorescent proteins at its N-terminus was expressed in COS-7 cells using the mammalian expression constructs pVenus-gephyrin (1) and the derived plasmid pmEos2-gephyrin (2), as well as the lentivirus replicons FU-mEos4b-gephyrin (3) and the newly generated variant FU-Dendra2-gephyrin (this study). Lentivirus was produced in HEK293 cells using the plasmids FU-Dendra2-gephyrin and FU-SP-myc-Dendra2-GlyR$\alpha$1 (4) as described previously (5).

#### COS-7 cell culture and photoconversion experiments

COS-7 cells were cultured on 18 mm diameter glass coverslips (VWR) in DMEM medium containing glutamax, 10% fetal bovine serum, 50 U/ml penicillin, and 50 µg/ml streptomycin at 37°C and 5% $CO_2$. When they had reached approximately 25% of confluence, cells were transfected with 0.5 µg plasmid DNA using FuGENE 6. After 24 hours of expression the cells were used for live imaging in Tyrode solution, or fixed at 37°C with 4% paraformaldehyde (PFA) in PBS for 15 minutes, rinsed, and imaged in PBS.

To compare the photo-physical properties of the fluorescent proteins expressed in COS-7 cells, the following image sequence was initiated: three wide-field fluorescence images were taken in the green channel (560/25 nm, 200 ms), followed by a focussed 405 nm laser beam (1 s, 5% of maximum power) and three more images in the green channel. Using FIJI software, the three images before the 405 nm pulse and the three last images were averaged and then divided to produce ratiometric images ($I_{post}$ / $I_{pre}$). The mean relative pixel intensities within three concentric rings (0.5-2.5 µm, 2.5-5 µm, 5-7.5 µm radius) around the bleached central spot (1 µm diameter) were then calculated and averaged for each fluorescent construct and experimental condition (n = 12 cells per condition).

#### Immunocytochemistry (ICC)

Cultured spinal cord neurons were fixed in PBS containing 4% paraformaldehyde (PFA) for 10 minutes at 37°C, permeabilised with 0.1% Triton X100 in PBS for 10 minutes, and blocked with 3% bovine serum albumin (BSA) in PBS for one hour. Primary antibodies (rabbit anti-GlyRα1, custom-made, 1:1000 dilution; mouse anti-VIAAT, 1:500) and secondary antibodies (Alexa Fluor 647-conjugated donkey anti-rabbit and donkey anti-mouse, Cy3-conjugated goat anti-mouse IgG, Jackson ImmunoResearch, 1:1000) were applied in blocking solution for one hour. The labelled neurons were imaged in PBS on a microscope setup similar to the one used for FRAP/FDAP, except for the illumination source (Intensilight, Nikon), the choice of emission filters (Semrock single-band bandpass filters, 525/30 nm for Dendra2, 607/36 nm for Cy3, 684/24 nm for A647), and the absence of the Ti-FRAP module. For the visualisation of crosslinked GlyRs, the receptors were immuno-immobilised with anti-GlyRα1 and A647-conjugated anti-rabbit IgG in living neurons (see Methods on antibody crosslinking), followed by fixation and ICC of the vesicular inhibitory amino acid transporter (VIAAT).

**Data curation of supplementary FRAP/FDAP data**

Out of the acquired primary data, the following recordings were removed in the supplementary figures: Fig. S2, no film removed. Fig. S3, one film removed for excess initial fluorescence, one film removed for lack of photoconversion signal. Fig. S5, CTRL: one film removed for lack of photoconversion signal, IMMO: no film removed.

### <u>References</u>

### Supplementary Figures

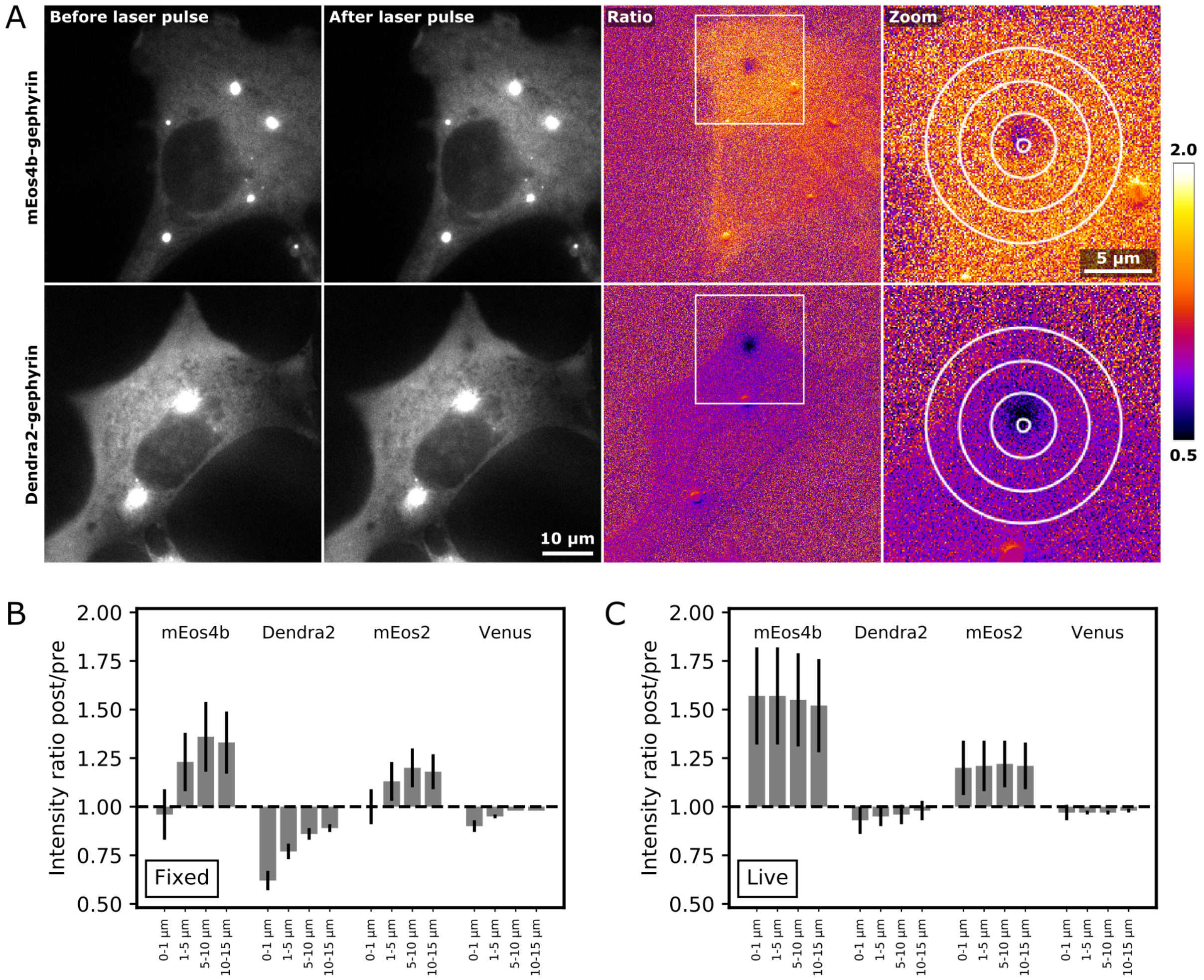


**Figure S1. Photo-physical properties of photoconvertible proteins**

(A) Gephyrin was tagged at its N-terminus with different fluorescent proteins (Dendra2, mEos2, mEos4b, Venus). Upon transfection in COS-7 cells, all constructs accumulated in large intracellular clusters, characteristic of overexpressed gephyrin (6). Fixed COS-7 cells expressing mEos4b-gephyrin (top panels) and Dendra2-gephyrin (lower panels) were exposed to a focussed 405 nm laser pulse. Images from left to right: 1. before and 2. after exposure to the laser beam, 3. ratiometric image (false colour pixel intensities $I_{post}$ / $I_{pre}$, between 0.5 and 2 fold), 4. zoomed region with circular zones of 1 µm, 5 µm, 10 µm, and 15 µm in diameter around the focussed laser spot.

(B) Quantification of relative changes in intensity of fluorescent proteins by 405 nm illumination in fixed and live COS-7 cells. Columns represent average intensity ratios ($I_{post}$ / $I_{pre}$) within each circular region (mean ± SD, n = 12 per condition). Note the increased fluorescence intensity of mEos4b and mEos2 around the central spot, both in fixed and even more so in live cells. This photochromic effect is less apparent for Dendra2-gephyrin, where photoconversion and/or bleaching of the fluorescent proteins dominate. In live imaging, the relative intensity changes are even throughout the cell due to the high mobility of gephyrin and the slow switching of the dichroic mirror and image acquisition.

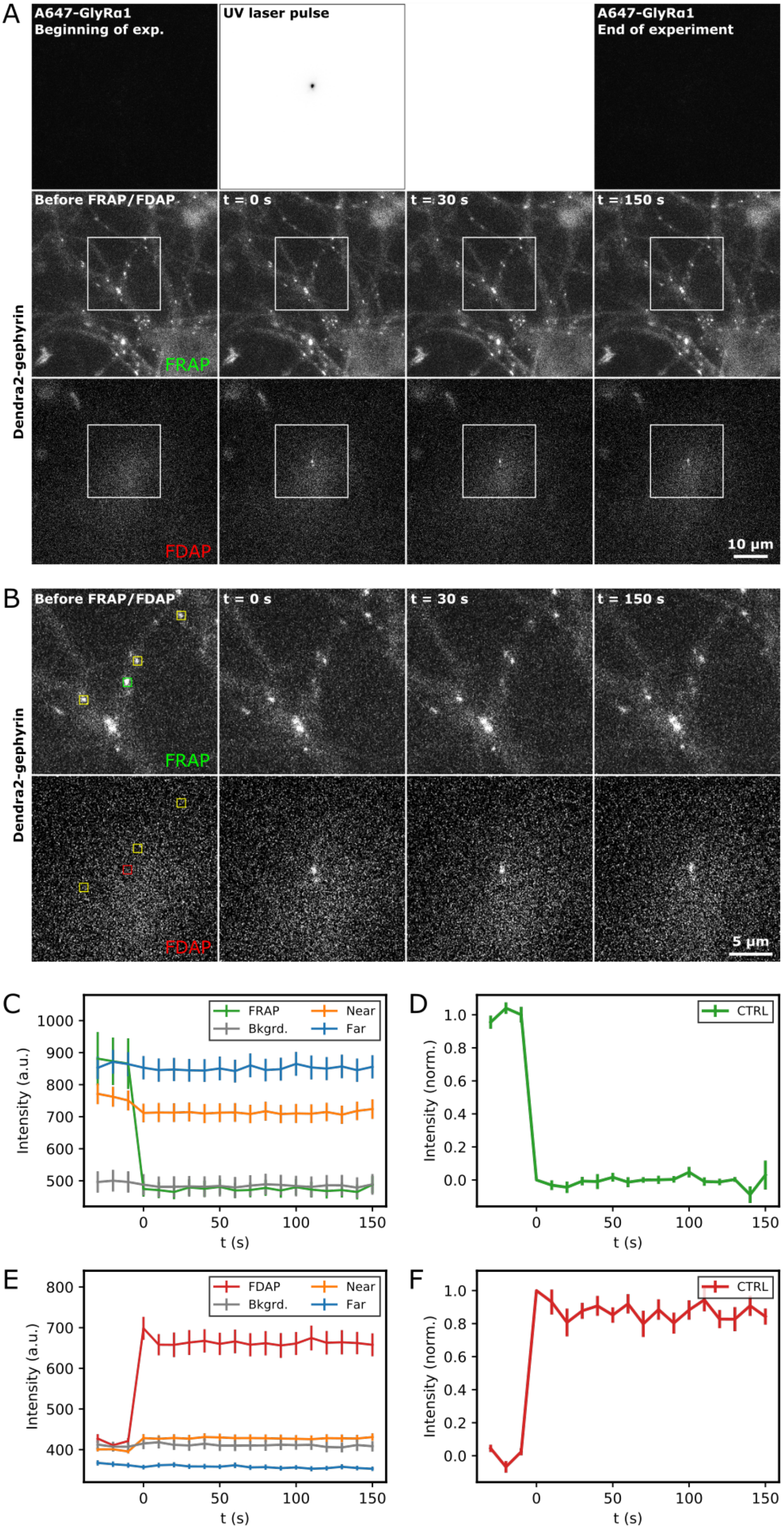
A
A647-GlyRα1
Beginning of exp.
UV laser pulse
A647-GlyRα1
End of experiment
Before FRAP/FDAP
t = 0 s
t = 30 s
t = 150 s
Dendra2-gephyrin
FRAP
FDAP
10 µm
B
Before FRAP/FDAP
t = 0 s
t = 30 s
t = 150 s
Dendra2-gephyrin
FRAP
FDAP
5 µm
C
FRAP
Near
Bkgrd.
Far
Intensity (a.u.)
t (s)
D
CTRL
Intensity (norm.)
t (s)
E
FDAP
Near
Bkgrd.
Far
Intensity (a.u.)
t (s)
F
CTRL
Intensity (norm.)
t (s)

**Figure S2. Methodology: fixed FRAP/FDAP recordings**

(A) FRAP/FDAP acquisition sequence in fixed neurons expressing Dendra2-gephyrin. Images in the far red channel were acquired at the beginning and at the end of the experiment to visualize A647-labelled, immuno-immobilised endogenous GlyRs (no signals under control conditions in the absence of receptor crosslinking). A 405 nm laser pulse was applied at time zero to photoconvert green Dendra2 fluorophores into the red state. Time-lapse images were recorded at 10 second intervals in the green (middle row) and the red channels (bottom row) before and after photoconversion. From left to right: 1. before exposure, 2. t = 0 (frame 1, after exposure to the focussed 405 nm laser beam), 3. t = 30 s (frame 4), 4. t = 150 s (frame 16).

(B) Zoomed regions of the time-lapse images in the green and red channels shown in A. The mean fluorescence intensity at synaptic puncta was measured in square areas of 9 x 9 pixels. The cluster in the centre of the image was used for FRAP/FDAP (shown in green and red, respectively); the yellow areas indicate near control points; the far control points are outside the shown region since they were selected at the edge of the field of view and as far as possible from the centre.

(C-F) Quantification of the mean fluorescence intensity of Dendra2-gephyrin puncta in fixed neurons (arbitrary units a.u., mean ± SEM, n = 22 fields of view). The green (C) and red traces (E) show the intensity of the central punctum exposed to the 405 nm laser beam in FRAP and FDAP mode, respectively. The orange and blue traces represent the intensity of the near and far control puncta, while the grey traces represent the diffuse signals of the Dendra2-gephyrin expressing neurons. The raw intensity data were normalised as described in the Methods section in both the green (D) and the red channels (F).

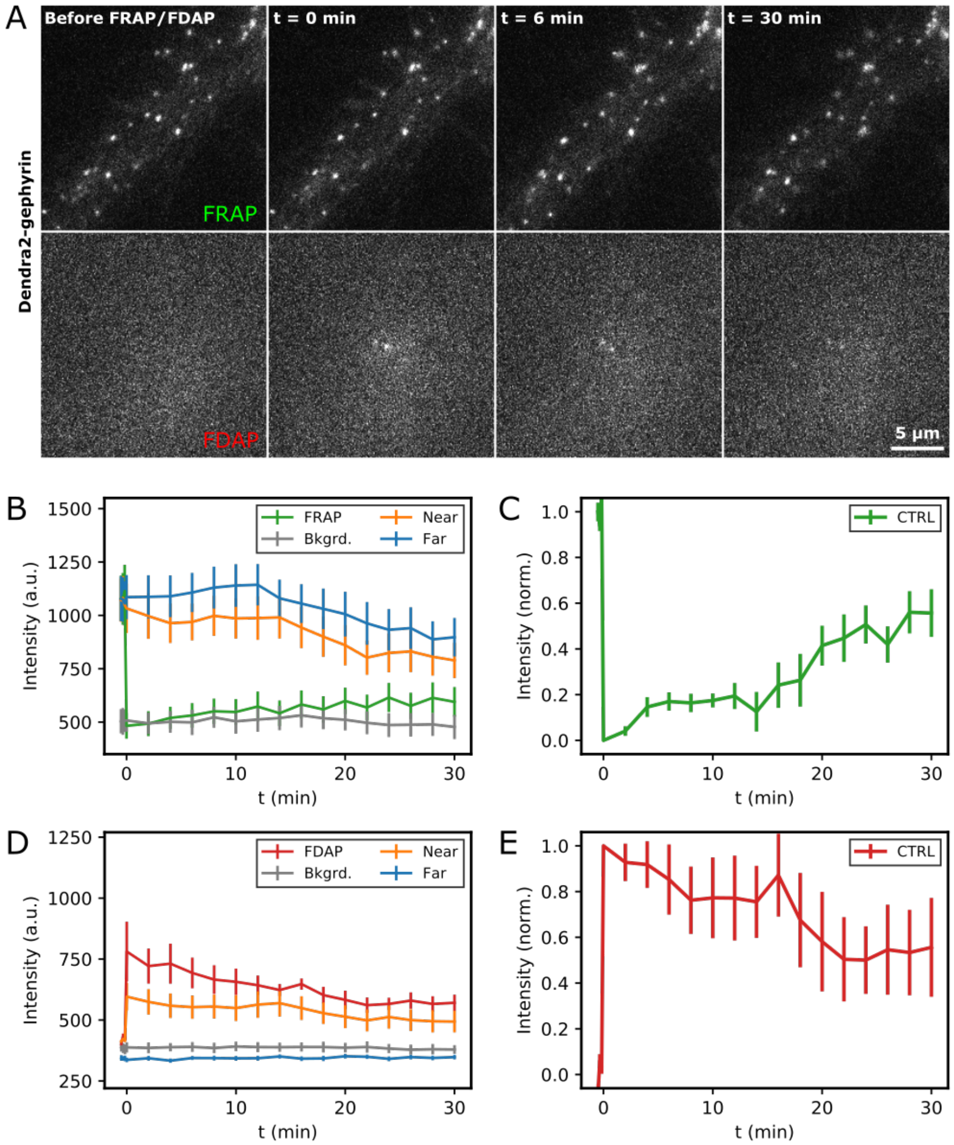


**Figure S3. Live FRAP/FDAP of Dendra2-gephyrin (preliminary experiments)**

(A) FRAP/FDAP time-lapse imaging of Dendra2-gephyrin with a 30 s acquisition frequency in living spinal cord neurons. Images from left to right 1. before exposure, 2. t = 0 (after exposure to a focussed 405 nm laser beam), 3. t = 6 min (frame 4), 4. t = 30 min (frame 16).

(B-E) Quantification of the mean Dendra2-gephyrin fluorescence intensity in the green (B, FRAP) and the red channel (D, FDAP) in living neurons (pilot experiments, mean a.u. ± SEM, n = 6 cells). Raw intensity data were normalised as described in the Methods section in the two channels (C,E).

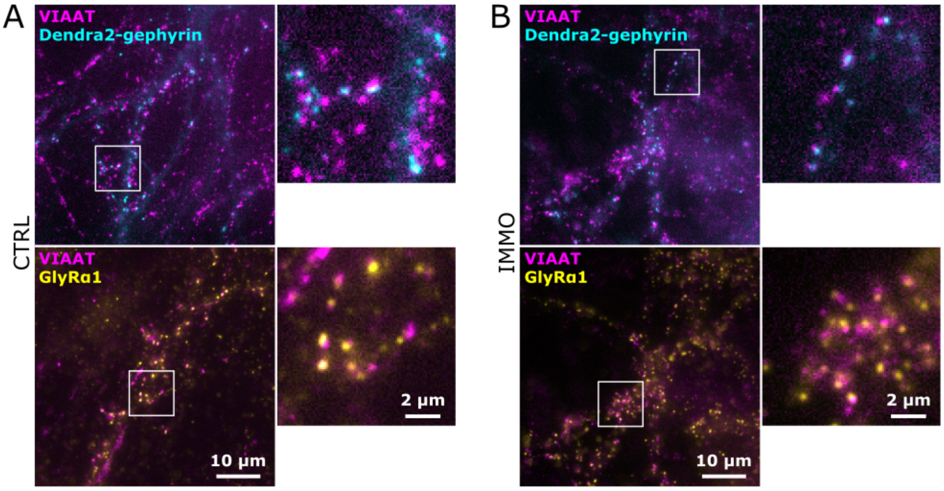


**Figure S4. Synaptic localisation of Dendra2-gephyrin and GlyRs**

(A) Immunocytochemistry of fixed spinal cord neurons (control condition). Top: Dendra2-gephyrin puncta (cyan) co-localise with presynaptic VIAAT (magenta). Bottom: ICC of endogenous anti-GlyRα1 (yellow) and VIAAT (magenta) at inhibitory synapses.

(B) Neurons were crosslinked with GlyRα1 antibody (immuno-immobilised condition) prior to fixation and immunolabelling. Top: Dendra2-gephyrin expression (cyan) and ICC for VIAAT (magenta). Bottom: Labelling of GlyRα1 (yellow) and VIAAT (magenta). GlyRα1 crosslinking did not have an obvious effect on the synaptic localisation of GlyRs in non-infected as well as Dendra2-gephyrin expressing neurons. Note that the top and bottom panels are two-colour overlays of the same field of view.

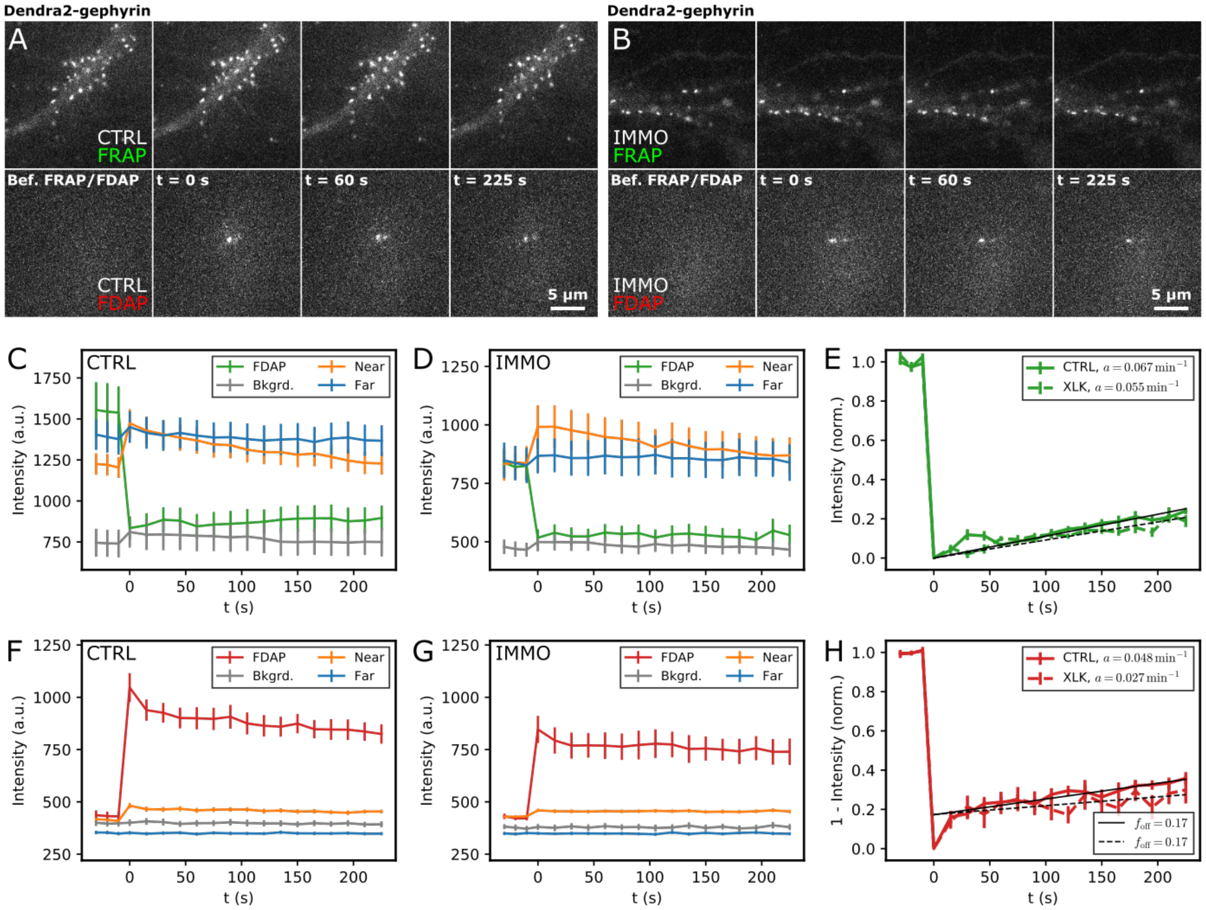


**Figure S5. High frequency recordings of Dendra2-gephyrin dynamics**

(A,B) FRAP/FDAP of Dendra2-gephyrin in spinal cord neurons under control conditions (A) and after GlyR immuno-immobilisation (B). Time-lapse imaging was done with a 15 s frequency over 4 min. Images from left to right 1. before exposure, 2. t = 0 (after the 405 nm laser pulse), 3. t = 60 s (frame 5), 4. t = 225 s (frame 16).

(C-H) Quantification of the mean fluorescence intensity of Dendra2-gephyrin in FRAP (C,D) and FDAP mode (F,G) under control conditions and after GlyR immuno-immobilisation (arbitrary units a.u., mean ± SEM, $n_{CTRL}$ = 15, $n_{IMMO}$ = 14 fields of view). (E,H) Normalised data.
