## Supplementary material for "Reciprocal stabilisation of glycine receptors and gephyrin scaffold proteins at inhibitory synapses": SI Text

### I. DESCRIPTION OF THE MODEL

Here, we propose and discuss a simple model of receptor-scaffold interactions. Our aim is to quantitatively reproduce the key observations from the immuno-immobilization experiments, and thus to propose a mechanistic interpretation of our experimental results. We intend the model to be a phenomenological description rather than a complete and microscopically detailed account of receptor and scaffold dynamics at the synapse.

For simplicity, we only consider three "species" and/or molecular states at the synapse: (i) loosely interacting scaffold proteins $s$, (ii) receptors $r$ that are diffusing and/or transiently attached to these scaffold proteins, and (iii) a population of more strongly linked receptor-scaffold complexes $c$. The dynamics of receptor and scaffold populations at the synapse arise from transitions between the states $s$, $r$ and $c$ and incoming as well as outgoing protein fluxes. At the stationary state, all these fluxes are balanced which allows us to determine the stationary values of the three considered populations.

#### A. Basic equations

A sketch of the proposed dynamics is shown in Fig. 5A in the main manuscript. We assume that loosely attached scaffolds and receptors (which may or may not already be bound) can bind and form, or join, with a rate $k_{\mathrm{b}}$ a more stable component $c$ of strongly interacting scaffold and receptors, while the inverse reaction occurs at a rate $k_{\mathrm{u}}$. Receptors that are not strongly interacting (i.e., that do not belong to $c$) leave the synapse laterally alone at a rate $j_{\mathrm{off}}$ or attached to scaffold proteins at a rate $g_{\mathrm{off}}$; they arrive at the synapse with a constant flux $J_{\mathrm{on}}$ without and $G_{\mathrm{on}}$ with scaffold protein attached. Eventually, scaffold proteins that are not strongly interacting (i.e., that do not belong to $c$) are desorbed into the cytoplasm at a rate $k_{\mathrm{off}}$ and leave the synapse together with receptors with the flux proportional to $g_{\mathrm{off}}$. They arrive by lateral diffusion with receptors with the flux $G_{\mathrm{on}}$ and are additionally recruited from the cytoplasm with a flux $K_{\mathrm{on}}$.

We introduce one additional parameter $\alpha$ that describes the stoichiometry of receptor to scaffold proteins in the strongly complexed state: For every scaffold protein added to the highly aggregated state, there are on average $\alpha$ receptors added, i.e., $s + \alpha r \to c$ with rate $r_{\mathrm{b}}$, and $c \to s + \alpha r$ with rate $k_{\mathrm{u}}$. Having in mind that synaptic gephyrin seems to occur almost exclusively in trimeric form, we consider that the "scaffold proteins" $s$ in the model correspond to gephyrin trimers. Based furthermore on evidence that GlyRs have two gephyrin binding sites that can potentially crosslink scaffolds by binding two different gephyrin trimers, we assume that in state $c$ the ratio of receptors to gephyrin trimers is about 3/2 on average, and we will consider $\alpha = 1.5$ (for an analysis of the influence of the value of $\alpha$ on the modeling outcomes, see appendix A). The total amount $R$ of receptors at the synapse is then given by

$$R = r + \alpha c \ , \tag{1}$$

while the total amount $S$ of scaffolds at the synapse is given by

$$S = s + c \ . \tag{2}$$

In other words, we count complexes $c$ by the number of involved scaffold proteins.

The equations that govern the dynamics of the different states $r$, $s$, and $c$ read as follows:

$$\frac{dr}{dt} = \alpha k_{\mathrm{u}} c - \alpha k_{\mathrm{b}} r s - (j_{\mathrm{off}} + g_{\mathrm{off}}) r + J_{\mathrm{on}} + G_{\mathrm{on}}, \tag{3}$$

$$\frac{ds}{dt} = k_{\mathrm{u}} c - k_{\mathrm{b}} r s - k_{\mathrm{off}} s - g_{\mathrm{off}} r + K_{\mathrm{on}} + G_{\mathrm{on}}, \tag{4}$$

$$\frac{dc}{dt} = -k_{\mathrm{u}} c + k_{\mathrm{b}} r s. \tag{5}$$

Note that we have chosen a simple second order reaction rate for the reactive fluxes $s + \alpha r \to c$, as we ignore the precise molecular reaction kinetics.

#### B. Stationary state

We can obtain the stationary states by setting the left-hand side (l.h.s.) of Eqs. (3)-(5) to zero. With $k_{\mathrm{u}} c = k_{\mathrm{b}} r s$ (Eq. (5)), we obtain the stationary value $r^*$ from Eq. (3), and subsequently $s^*$ and $c^*$ from Eqs. (4) and (5), respectively:

$$r^* = \frac{J_{\text{on}} + G_{\text{on}}}{j_{\text{off}} + g_{\text{off}}}, \tag{6}$$

$$s^* = \frac{K_{\text{on}} + G_{\text{on}} - g_{\text{off}} r^*}{k_{\text{off}}} = \frac{K_{\text{on}} + G_{\text{on}} - g_{\text{off}} \frac{J_{\text{on}} + G_{\text{on}}}{j_{\text{off}} + g_{\text{off}}}}{k_{\text{off}}}, \tag{7}$$

$$c^* = \frac{k_{\text{b}}}{k_{\text{u}}} r^* s^* = \frac{k_{\text{b}}}{k_{\text{u}}} \frac{J_{\text{on}} + G_{\text{on}}}{j_{\text{off}} + g_{\text{off}}} \frac{K_{\text{on}} + G_{\text{on}} - g_{\text{off}} \frac{J_{\text{on}} + G_{\text{on}}}{j_{\text{off}} + g_{\text{off}}}}{k_{\text{off}}}. \tag{8}$$

The total amounts of receptors and scaffolds in the stationary state are given by

$$R^* = \frac{J_{\text{on}} + G_{\text{on}}}{j_{\text{off}} + g_{\text{off}}} \left( 1 + \alpha \frac{k_{\text{b}}}{k_{\text{u}}} \frac{K_{\text{on}} + G_{\text{on}} - g_{\text{off}} \frac{J_{\text{on}} + G_{\text{on}}}{j_{\text{off}} + g_{\text{off}}}}{k_{\text{off}}} \right), \tag{9}$$

$$S^* = \left( 1 + \frac{k_{\text{b}}}{k_{\text{u}}} \frac{J_{\text{on}} + G_{\text{on}}}{j_{\text{off}} + g_{\text{off}}} \right) \frac{K_{\text{on}} + G_{\text{on}} - g_{\text{off}} \frac{J_{\text{on}} + G_{\text{on}}}{j_{\text{off}} + g_{\text{off}}}}{k_{\text{off}}}. \tag{10}$$

In the following, we choose to normalize all protein amounts by $S^*$, as well as the fluxes $J_{\text{on}}$, $G_{\text{on}}$, and $K_{\text{on}}$. To eliminate $S^*$ from the resulting equations for the rescaled amounts, we furthermore rescale $k_{\text{b}} \to S^* k_{\text{b}}$. Our normalization allows us to eliminate one parameter from the dynamic equations above. Using Eq. (10) and $S^* = 1$ in rescaled units, we can express the rescaled cytoplasmic scaffold influx $K_{\text{on}}$ as a function of all other parameters,

$$K_{\text{on}} = \frac{k_{\text{off}}}{1 + \frac{k_{\text{b}}}{k_{\text{u}}} \frac{J_{\text{on}} + G_{\text{on}}}{j_{\text{off}} + g_{\text{off}}}} - G_{\text{on}} + g_{\text{off}} \frac{J_{\text{on}} + G_{\text{on}}}{j_{\text{off}} + g_{\text{off}}}. \tag{11}$$

Note that we keep the original symbols for the rescaled quantities in order to keep the notation simple. In rescaled units, the stationary states then become

$$r^* = \frac{J_{\text{on}} + G_{\text{on}}}{j_{\text{off}} + g_{\text{off}}}\ , \tag{12}$$

$$s^* = \frac{1}{1 + \frac{k_{\text{b}}}{k_{\text{u}}} \frac{J_{\text{on}} + G_{\text{on}}}{j_{\text{off}} + g_{\text{off}}}}, \tag{13}$$

$$c^* = \frac{1}{1 + \frac{k_{\text{u}}}{k_{\text{b}}} \frac{j_{\text{off}} + g_{\text{off}}}{J_{\text{on}} + G_{\text{on}}}}. \tag{14}$$

For the rescaled total amount of receptors, this naturally implies

$$R^* = \frac{J_{\text{on}} + G_{\text{on}}}{j_{\text{off}} + g_{\text{off}}} + \frac{\alpha}{1 + \frac{k_{\text{u}}}{k_{\text{b}}} \frac{j_{\text{off}} + g_{\text{off}}}{J_{\text{on}} + G_{\text{on}}}}. \tag{15}$$

#### C. FRAP/FDAP dynamics

In the model, the FRAP and FDAP in the stationary state are equivalent as all fluxes are perfectly balanced. For simplicity, we will write the equations describing the loss of photoconverted protein, which is more easily captured mathematically.

#### 1. Receptor FDAP

In order to model a FDAP experiment for the receptor, we consider that at time $t = 0$, all receptors present at the synapse are photoconverted and followed in time, while incoming fluxes are not contributing to the visible population. At the same time, the scaffold population is not affected. We will therefore consider only the dynamics of visible populations $\hat{r}(t)$ and $\hat{c}(t)$, the dynamics of which are governed by the equations

$$\frac{d\hat{r}}{dt} = \alpha k_{\mathrm{u}}\hat{c} - \alpha k_{\mathrm{b}}\hat{r}s^* - (j_{\mathrm{off}} + g_{\mathrm{off}})\hat{r}, \tag{16}$$

$$\frac{d\hat{c}}{dt} = -k_{\mathrm{u}}\hat{c} + k_{\mathrm{b}}\hat{r}s^*. \tag{17}$$

Because of a possible transitioning via the state $c$, the decay of the visible receptor population does not follow a simple exponential decay with a characteristic timescate $1/(j_{\mathrm{off}}+g_{\mathrm{off}})$ but follows a bi-exponential dynamics that we can determine as follows. If we express the above equation as

$$\frac{d}{dt}\begin{pmatrix}\hat{r}\\ \hat{c}\end{pmatrix} = -M^r\begin{pmatrix}\hat{r}\\ \hat{c}\end{pmatrix}\quad , \qquad M^r = \begin{pmatrix}\alpha k_{\mathrm{b}}s^* + j_{\mathrm{off}} + g_{\mathrm{off}} & -\alpha k_{\mathrm{u}}\\ -k_{\mathrm{b}}s^* & k_{\mathrm{u}}\end{pmatrix} \tag{18}$$

we can identify the matrix $M^r$ that governs the linear dynamics of $\hat{r}$ and $\hat{c}$. The solution of the dynamics is given by the eigenvectors $(r_1, c_1)^T, (r_2, c_2)^T$ and eigenvalues $k_1, k_2$ of $M$ such that $\hat{r}(t) = ar_1e^{-k_1t} + br_2e^{-k_2t}$ and $\hat{c}(t) = ac_1e^{-k_1t} + bc_2e^{-k_2t}$. (The constants $a$ and $b$ are chosen such that the initial conditions $\hat{r}(0) = r^*$ and $\hat{c}(0) = c^*$ are satisfied.) The two characteristic decay rates are relatively straightforward to obtain, and one eventually gets

$$k^r_{1,2} = \frac{\alpha k_{\mathrm{b}}s^* + j_{\mathrm{off}} + g_{\mathrm{off}} + k_{\mathrm{u}}}{2} \pm \sqrt{\frac{(\alpha k_{\mathrm{b}}s^* + j_{\mathrm{off}} + g_{\mathrm{off}} - k_{\mathrm{u}})^2}{4} + \alpha k_{\mathrm{u}}k_{\mathrm{b}}s^*}. \tag{19}$$

For comparison with the experimental FRAP/FDAP data, we assume that the fluorescence intensity is proportional to the amount of visible receptors and compute the normalized trace

$$I_r(t) = \frac{\hat{r}(t) + \alpha\hat{c}(t)}{\hat{r}(0) + \alpha\hat{c}(0)} = \frac{\hat{r}(t) + \alpha\hat{c}(t)}{r^* + \alpha c^*}. \tag{20}$$

#### 2. Scaffold FDAP

In the case of photoconverted scaffold proteins, we have to track the scaffold populations while considering the receptor concentration to remain constant. Similarly to the calculation above, we can describe the coupled dynamics of photoconverted populations $s$ and $c$ by

$$\frac{d\hat{s}}{dt} = k_\mathrm{u}\hat{c} - k_\mathrm{b} r^* \hat{s} - \left(k_\mathrm{off} + g_\mathrm{off}\frac{r^*}{s^*}\right)\hat{s}, \tag{21}$$

$$\frac{d\hat{c}}{dt} = -k_\mathrm{u}\hat{c} + k_\mathrm{b} r^* \hat{s}, \tag{22}$$

where the factor $r^*/s^*$ accounts for the fact that the flux of laterally exiting receptor-scaffold complexes is given by $g_\mathrm{off} r^*$ in the stationary state.

We can again write the evolution of $\hat{s}$ and $\hat{c}$ in compact form,

$$\frac{d}{dt}\begin{pmatrix}\hat{s}\\ \hat{c}\end{pmatrix} = -M^s \begin{pmatrix}\hat{s}\\ \hat{c}\end{pmatrix}, \qquad M^s = \begin{pmatrix} k_\mathrm{b} r^* + k_\mathrm{off} + g_\mathrm{off}\frac{r^*}{s^*} & -k_\mathrm{u} \\ -k_\mathrm{b} r^* & k_\mathrm{u}\end{pmatrix} \tag{23}$$

and subsequently obtain the characteristic decay rates

$$k^s_{1,2} = \frac{k_\mathrm{b} r^* + k_\mathrm{off} + g_\mathrm{off}\frac{r^*}{s^*} + k_\mathrm{u}}{2} \pm \sqrt{\frac{(k_\mathrm{b} r^* + k_\mathrm{off} + g_\mathrm{off}\frac{r^*}{s^*} - k_\mathrm{u})^2}{4} + k_\mathrm{u} k_\mathrm{b} r^*}. \tag{24}$$

For comparison with the experimental FRAP/FDAP data, we assume that the fluorescence intensity is proportional to the amount of visible scaffolds and compute the normalized trace

$$I_s(t) = \hat{s}(t) + \hat{c}(t). \tag{25}$$

Importantly, our analysis demonstrates that the intra-synaptic exchange between loosely bound and stable pools gives rise to a bi-exponential FRAP/FDAP relaxation dynamics with two characteristic timescales both for receptors and for scaffolds. On intermediate timescales, such a bi-exponential dynamics may ressemble a single exponential relaxation with a finite stable fraction. In contrast, if no such intra-synaptic exchange took place e.g. because no stable component existed, all FRAP/FDAP relaxation dynamics would be governed by a single characteristic timescale and no apparent stable fraction could be observed.

#### D. Dwell times

We can as well calculate the distribution of dwell times of receptors and scaffolds after entry at the synapse. The probabilities for a newly entered receptor to be in state $r$ or $c$ evolve according to equations analogous to Eqs. (16) and (17), that is,

$$\frac{dp_r}{dt} = \alpha k_\mathrm{u} p_c - \alpha k_\mathrm{b} p_r s^* - (j_\mathrm{off} + g_\mathrm{off}) p_r, \tag{26}$$

$$\frac{dp_c}{dt} = -k_\mathrm{u} p_c + k_\mathrm{b} p_r s^*. \tag{27}$$

The probability $P_r(t) = p_r(t) + p_c(t)$ for a receptor to be still be at the synapse at time $t$ can be obtained along the same lines as the FDAP signal $I_r(t)$ above, using the initial conditions $p_r(0) = 1$ and $p_c(t) = 0$. The cumulative dwell time distribution of receptors is then simply given by $1 - P_r(t)$.

For scaffolds, we proceed analogously using the equations

$$\frac{dp_s}{dt} = k_{\mathrm{u}} p_c - k_{\mathrm{b}} r^* p_s - \left( k_{\mathrm{off}} + g_{\mathrm{off}} \frac{r^*}{s^*} \right) p_s, \tag{28}$$

$$\frac{dp_c}{dt} = -k_{\mathrm{u}} p_c + k_{\mathrm{b}} r^* p_s \tag{29}$$

with initial conditions $p_s(0) = 1$ and $p_c(t) = 0$. The cumulative dwell time distribution is accordingly given by $1 - p_s(t) - p_c(t)$.

### II. THE EFFECT OF IMMUNO-IMMOBILIZATION

In the immuno-immobilized condition (IMMO in the main manuscript), we assume that receptors are immobilized in- and outside of the synapse and can neither enter not exit the synaptic domain. As a consequence, total receptor numbers at the synapse remain constant after immobilization, although their relative proportion in the different states may change. We account for this latter possibility by considering that a fraction $f$ of loosely bound receptors $r^*$ eventually ends up in the highly crosslinked state $c$ together with a corresponding fraction of loosely bound scaffolds of state $s$. We thus stipulate

$$r^{\times} = (1 - f)\, r^*, \tag{30}$$

$$c^{\times} = c^* + f r^*/\alpha, \tag{31}$$

where the superscript $\times$ denotes the immobilized condition.

Since all fluxes involving receptors are suppressed, the effective dynamics of the remaining synaptic "species" $s$ after immobilization is then simply described by

$$\frac{ds}{dt} = -k_{\mathrm{off}} s + K_{\mathrm{on}}, \tag{32}$$

with $K_{\mathrm{on}}$ unchanged. In the stationary state,

$$s^{\times} = K_{\mathrm{on}}/k_{\mathrm{off}}. \tag{33}$$

Our experimental observation that the synaptic size is not affected by receptor immobilization implies that $S^\times = s^\times + c^\times = 1$, and thus $s^\times = s^* - fr^*/\alpha$. From this follows (with Eqs. (11) and (13)) an additional constraint on the parameters, namely that

$$G_{\mathrm{on}} = \frac{g_{\mathrm{off}} + \frac{k_{\mathrm{off}} f}{\alpha}}{j_{\mathrm{off}} - \frac{k_{\mathrm{off}} f}{\alpha}} J_{\mathrm{on}}. \tag{34}$$

In the case of $f = 0$, we have $G_{\mathrm{on}} = \frac{g_{\mathrm{off}}}{j_{\mathrm{off}}} J_{\mathrm{on}}$ and find that all fluxes are individually balanced in the stationary state: $G_{\mathrm{on}} = g_{\mathrm{off}} r^*$, $J_{\mathrm{on}} = j_{\mathrm{off}} r^*$, and $K_{\mathrm{on}} = k_{\mathrm{off}} s^*$. This implies that the system is in thermodynamic equilibrium in this case.

Because the strongly connected component $c$ does not turn over in the immobilized condition, we can now easily determine the FDAP time course of the scaffold protein. From Eq. (32), we see that in the stationary state, the loosely aggregated scaffold protein population $s$ simply observes an exponential decay with a single time constant $1/k_{\mathrm{off}}$ towards the stable fraction $c^\times$,

$$I_s^\times(t) = s^\times e^{-t k_{\mathrm{off}}} + c^\times. \tag{35}$$

We furthermore note that if no stable component $c$ existed, the scaffold FDAP relaxation dynamics in the immobilized condition would relax toward 0 as no stable fraction $c^\times$ existed.

### III. FIT RESULTS

Using the above analytical results, we can now try to find optimal parameters for which the model best approximates the experimental data. More precisely, we fit the combined FRAP/FDAP data for GlyRs in the control condition and gephyrin in control and immuno-immobilized conditions with our model using a least-squares fit. This allows us to determine the remaining 6 free parameters of our model $(k_{\mathrm{off}}, j_{\mathrm{off}}, g_{\mathrm{off}}, J_{\mathrm{on}}, k_{\mathrm{u}}, k_{\mathrm{b}})$ as a function of the value of $f$; parameters $K_{\mathrm{on}}$ and $G_{\mathrm{on}}$ follow by Eqs. (11) and (34).

A summary of our fit results is presented in Fig. 5B-F of the main manuscript. We show here the best fit parameter values with 95% confidence intervals as reported by the fit routine in Fig. I. The corresponding values for $r^*$, $s^*$, and $c^*$ the stationary state are shown in Fig. II, and the corresponding fluxes in Fig. III.

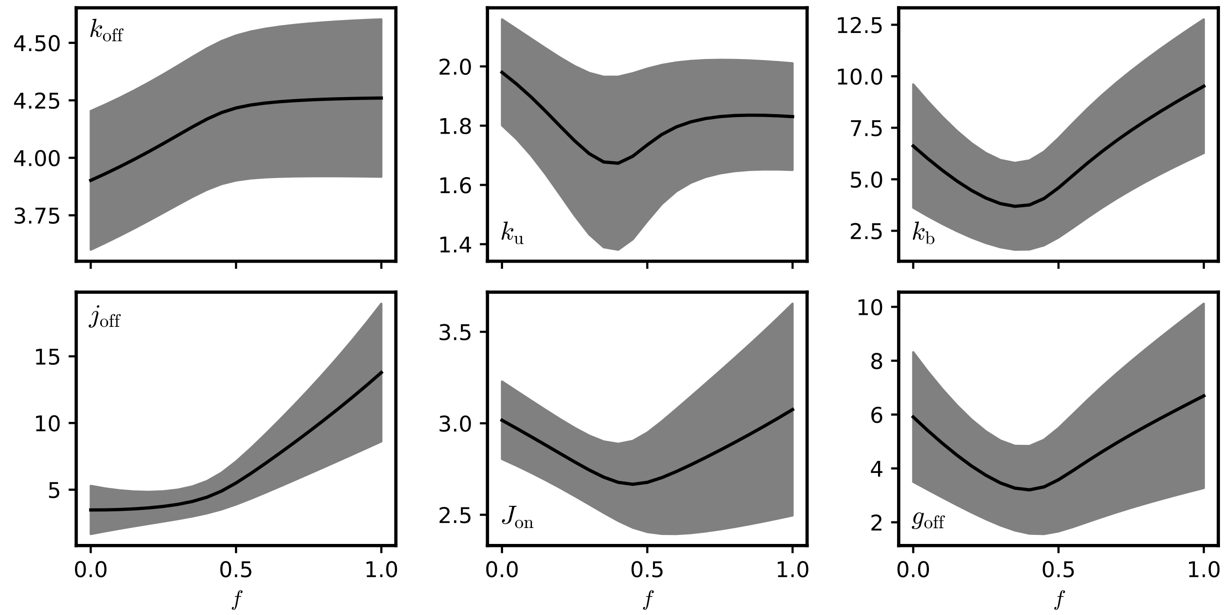


FIG. I. Variation of the best fit parameters with fraction $f$ of receptors converted into the tightly bound state $c$ after immobilization. All units are $\mathrm{h}^{-1}$; note that in our model all fluxes are rescaled by the total amount of scaffolds.

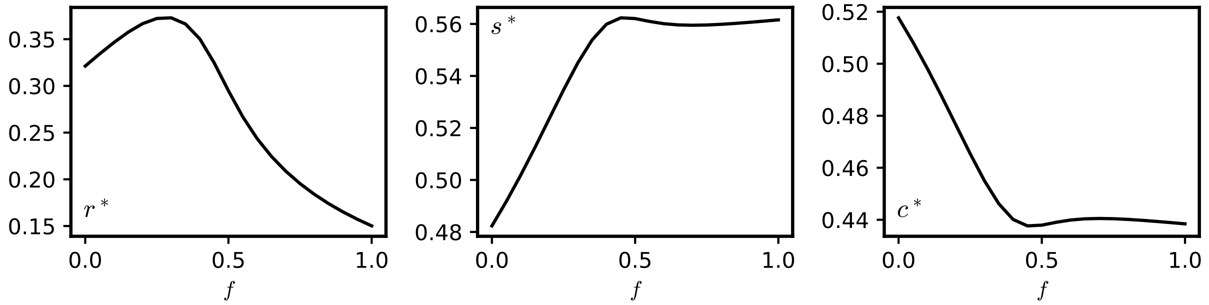


FIG. II. Fit results for the receptor and scaffold states $r^*$, $s^*$, and $c^*$ as a function of the fraction $f$ of receptors converted into the tightly bound state $c$ after immobilization.

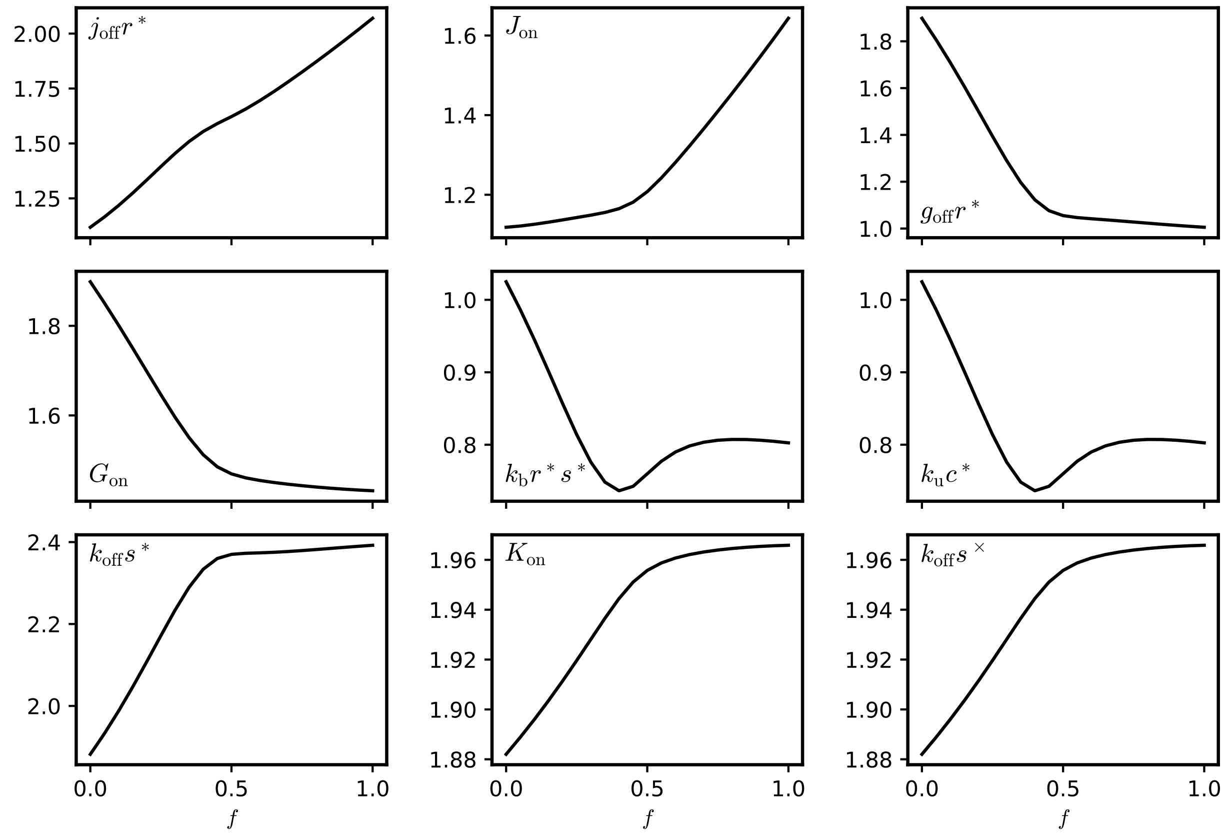


FIG. III. Fit results for the fluxes as a function of the fraction $f$ of receptors converted into the tightly bound state $c$ after immobilization.

#### Appendix A: Influence of the value of $\alpha$ on the fit results

In this work, we assume a fixed stoichiometry between receptors and scaffolds in state $c$. Motivated by the structural properties of glycine receptors and gephyrin trimers, we considered this stoichiometry to be $\alpha = 1.5$. However, we might ask how a different value of $\alpha$ impacts the fit results. The influence of the value of $\alpha$ on the fit parameters is shown in Fig. IV, where for simplicity we restricted ourselves to the case $f = 0$. All values are rescaled with respect to the reference $\alpha = 1.5$. While $k_{\mathrm{off}}$ and $k_{\mathrm{u}}$ do not change significantly, all other parameters do vary with $\alpha$, albeit rather slightly.

The effect on the states $r^*$, $s^*$, and $c^*$ in the control condition is shown in Fig. V. Interestingly, $s^*$ and $c^*$ remain constant throughout the whole range of values explored. The amount of loosely bound receptors seems to scale linearly with $\alpha$ as can be expected from the scaling of the receptor influx $J_{\mathrm{on}}$ (Fig. IV). Note however that with $r^* \propto \alpha$, $c^* = \mathrm{const.}$, the total synaptic amount of receptors $R^* = r^* + \alpha c^*$ will also scale linearly with $\alpha$.

The dependence on $\alpha$ of the fluxes is shown in Fig. VI. Note that only the predicted receptor influx $J_{\mathrm{on}}$ and efflux $j_{\mathrm{off}} r^*$ are affected by considering a different value of $\alpha$; all other fluxes remain basically unchanged.

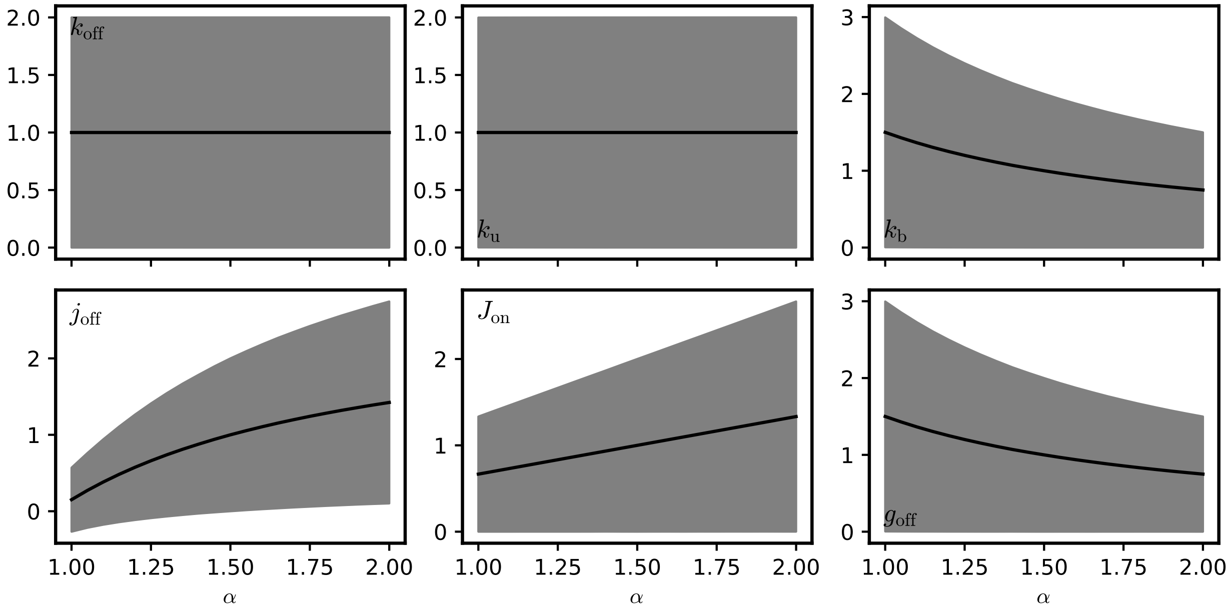


FIG. IV. Variation of the best fit parameters with $\alpha$. Parameter values are rescaled to their value for the case $\alpha = 1.5$ considered in the main manuscript.

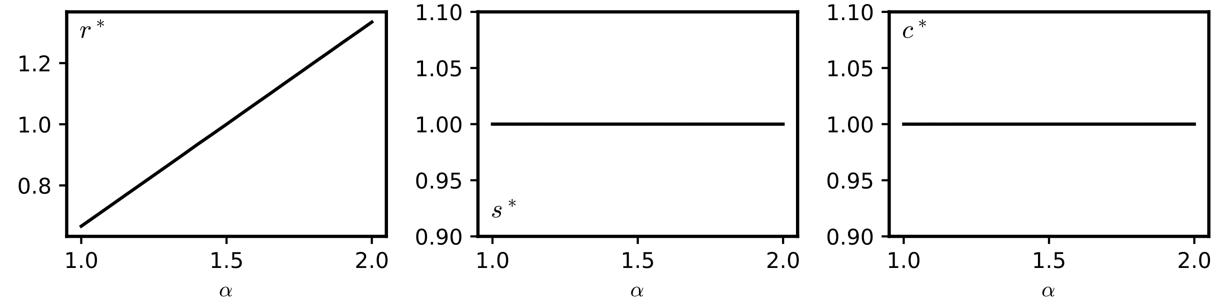


FIG. V. The fit results for the receptor and scaffold states $r^*$, $s^*$, and $c^s$ as a function of $\alpha$. Values are rescaled to their value for the case $\alpha = 1.5$ considered in the main manuscript.

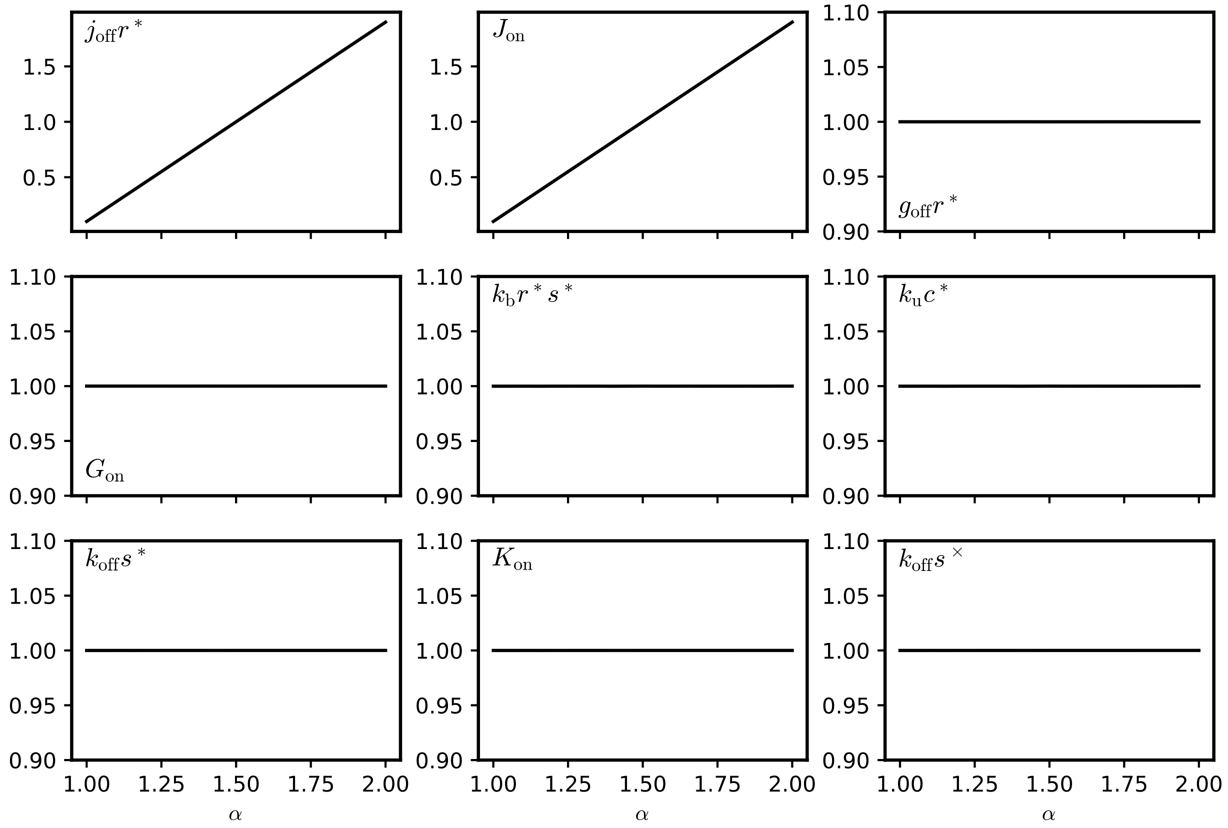


FIG. VI. Variation of the fluxes predicted by the model with $\alpha$. Fluxes are rescaled to their value for the case $\alpha = 1.5$ considered in the main manuscript.

**Appendix B: Comparison with a reduced model without entry or exit of laterally diffusing GlyR-gephyrin complexes ($G_{\rm on} = g_{\rm off} = 0$)**

In our model, we considered that receptors and scaffolds may enter and exit the synapse individually in exchange with extrasynaptic pools (pathways (1) and (4) in Fig. 5 of the main manuscript), and together in the form of receptor-scaffold complexes that are also present in the extrasynaptic membrane (pathway (2) in Fig. 5 of the main manuscript). Note that including the latter pathway in the model does not *per se* imply that it contributes to the synaptic dynamics: Only fitting our model to our experimental data allowed us to identify values for the parameters associated with all of these fluxes and conclude that all three pathways are involved in the exchange of receptors and scaffolds between synaptic and extrasynaptic pools.

We checked more specifically whether the recruitment of laterally diffusing receptor-scaffold complexes to the synapse and the loss of receptor-scaffold complexes to the extrasynaptic membrane needed to be taken into account to accurately fit the experimental data. To this end, we considered a reduced version of our model, where we imposed $G_{\rm on} = g_{\rm off} = 0$ in all of the model equations presented above. Instead of the six free parameters of the original model we are then left with only five effective parameters, as $G_{\rm on} = 0$ follows from $g_{\rm off} = 0$ and $f = 0$ with Eq. (34). (Note that if $G_{\rm on} = g_{\rm off} = 0$, the remaining receptor and scaffold fluxes need to be balanced individually, and the constant synaptic size then requires $f = 0$.)

The fit result for the reduced model is shown in Fig. VIIA. By visual inspection alone, the fit without entry and exit of receptor-scaffold complexes is markedly less accurate than than the fit with the full model (compare Fig. 5B,C of the main manuscript). In order to quantitatively compare the full and the reduced model, in consideration of the difference in model complexity as reflected by the different number of free parameters, we computed the Bayesian information criterion (BIC)[1] for the two models,

$$BIC = -2 \log \hat{L}(y; M, \hat{\theta}) + k \log n, \tag{B1}$$

where $\hat{L}(y; Mm\hat{\theta})$ is the Maximum-Likelihood of the observed data $y$ under the model $M$ (i.e. with best-fit parameters $\hat{\theta}$), $k$ is the number of parameters of the model, and $n$ is

[1] e.g. G. Claeskens and N. L. Hjort, Model Selection and Model Averaging, Cambridge University Press, Cambridge (2008)

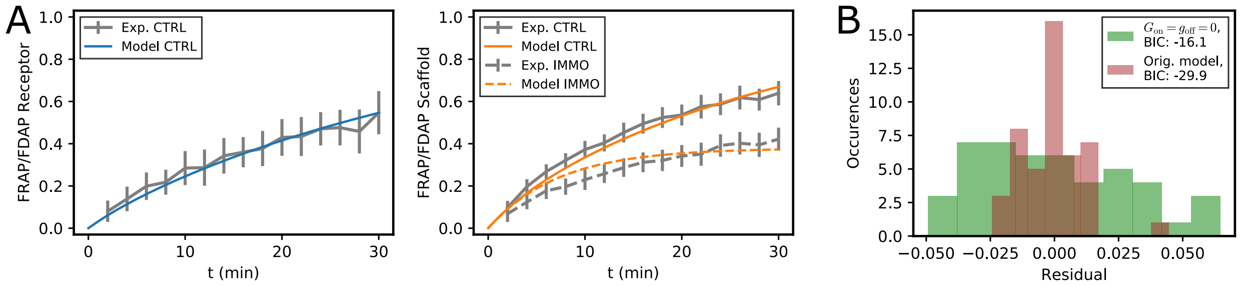


FIG. VII. Fit of the experimental data with a reduced model that does not account for lateral exchange of receptor-scaffold complexes. (A) Comparison of model curves for the best-fit parameters to experimental FRAP/FDAP curves for GlyR in the control condition (left) and gephyrin in control and immuno-immobilized conditions (right). (B) Distributions of tje residuals for the reduced and the full model (with $f = 0$).

the number of observations. The BIC formalizes the idea of a trade-off between “fitting the data better” (via the log-likelihood of the data) and “model simplicity” (via a penalty proportional to the number of model parameters). Under the assumption of independent, identically-distributed Gaussian noise on individual observations, the BIC becomes

$$BIC = n \log \frac{R}{n} + k \log n \tag{B2}$$

up to additive, model-independent constants. Here, $R = \sum_{i=1}^{n} (y_i - \hat{y}_i(M, \hat{\theta}))^2$ is the sum of squared residuals for the fitted model; the $y_i$ and $\hat{y}_i$ are the data points and the values predicted by the model, respectively. In our case, $n = 45$.

Histograms of the residuals for the original model (with $f = 0$) and for the reduced model are shown in Fig. VIIB, further indicating that the reduced model accounts less well for the experimental data. This is entirely corroborated by the difference of their BIC, with $BIC_{\text{reduced}} - BIC_{\text{full}} > 13$. For completeness, we end by stating that the BIC for the full model does hardly depend on the value of $f$, as all values lie in the range of (-30.18,-29.45).
